# Theory on the rate equation of Michaelis-Menten type single-substrate enzyme catalyzed reactions

**DOI:** 10.1101/143396

**Authors:** Rajamanickam Murugan

**Affiliations:** Department of Biotechnology, Indian Institute of Technology Madras Chennai, India

**Keywords:** Single-substrate enzyme, Michaelis-Menten kinetics, steady state approximation

## Abstract

Analytical solution to the Michaelis-Menten (MM) rate equations for single-substrate enzyme catalysed reaction is not known. Here we introduce an effective scaling scheme and identify the critical parameters which can completely characterize the entire dynamics of single substrate MM enzymes. Using this scaling framework, we reformulate the differential rate equations of MM enzymes over velocity-substrate, velocity-product, substrate-product and velocity-substrate-product spaces and obtain various approximations for both pre- and post-steady state dynamical regimes. Using this framework, under certain limiting conditions we successfully compute the timescales corresponding to steady state, pre- and post-steady states and also compute the approximate steady state values of velocity, substrate and product. We further define the dynamical efficiency of MM enzymes as the ratio between the reaction path length in the velocity-substrate-product space and the average reaction time required to convert the entire substrate into product. Here dynamical efficiency characterizes the phase-space dynamics and it would tell us how fast an enzyme can clear a harmful substrate from the environment. We finally perform a detailed error level analysis over various pre- and post-steady state approximations along with the already existing quasi steady state approximations and progress curve models and discuss the positive and negative points corresponding to various steady state and progress curve models.

## 1. Introduction

Enzymes are important biocatalysts which drive various reaction steps of all biological and biochemical pathways (Alberts, 2002; Stryer, 1988; Voet and Voet, 1995). The **M**ichaelis-**M**enten kinetic scheme (MMS) (Briggs and Haldane, 1925; Michaelis and Menten, 1913) is the fundamental mechanistic description of biological catalysis of enzyme reactions (Cornish-Bowden, 2015; Deichmann et al., 2014; Johnson and Goody, 2011). In this scheme (Fig. 1), enzyme molecule first binds with its substrate which is a reactant to form enzyme-substrate complex in reversible manner. We denote the forward and reverse rate constants of this first step as *k_1_* (M^−1^s^−1^) and *k_−1_* (s^−1^) respectively. Subsequently this enzyme-substrate complex will irreversibly dissociate into free enzyme and product with a rate of *k_2_* (s^−1^). Integral solution to the set of kinetic rate equations associated with the Michaelis-Menten scheme is not known. Several theoretical groups have tried to expand the integral solution of MMS in terms of ordinary (Murugan, 2002) and singular perturbation series (Murray, 2002). Perturbation expansions of the solution trajectory over slow manifolds have also been tried (Fraser, 2004; Roussel and Fraser, 2001). Singular perturbation expansions always yield a combination of inner and outer solutions corresponding to pre-steady state and post-steady state timescales which were then combined via proper matching at the temporal boundary layer (Dell’Acqua and Bersani, 2012; Dingee and Anton, 2008;Murray, 2002; Segel and Slemrod, 1989; Seshadri and Fritzsch, 1981; Vogt, 2013).

**FIGURE 1.**
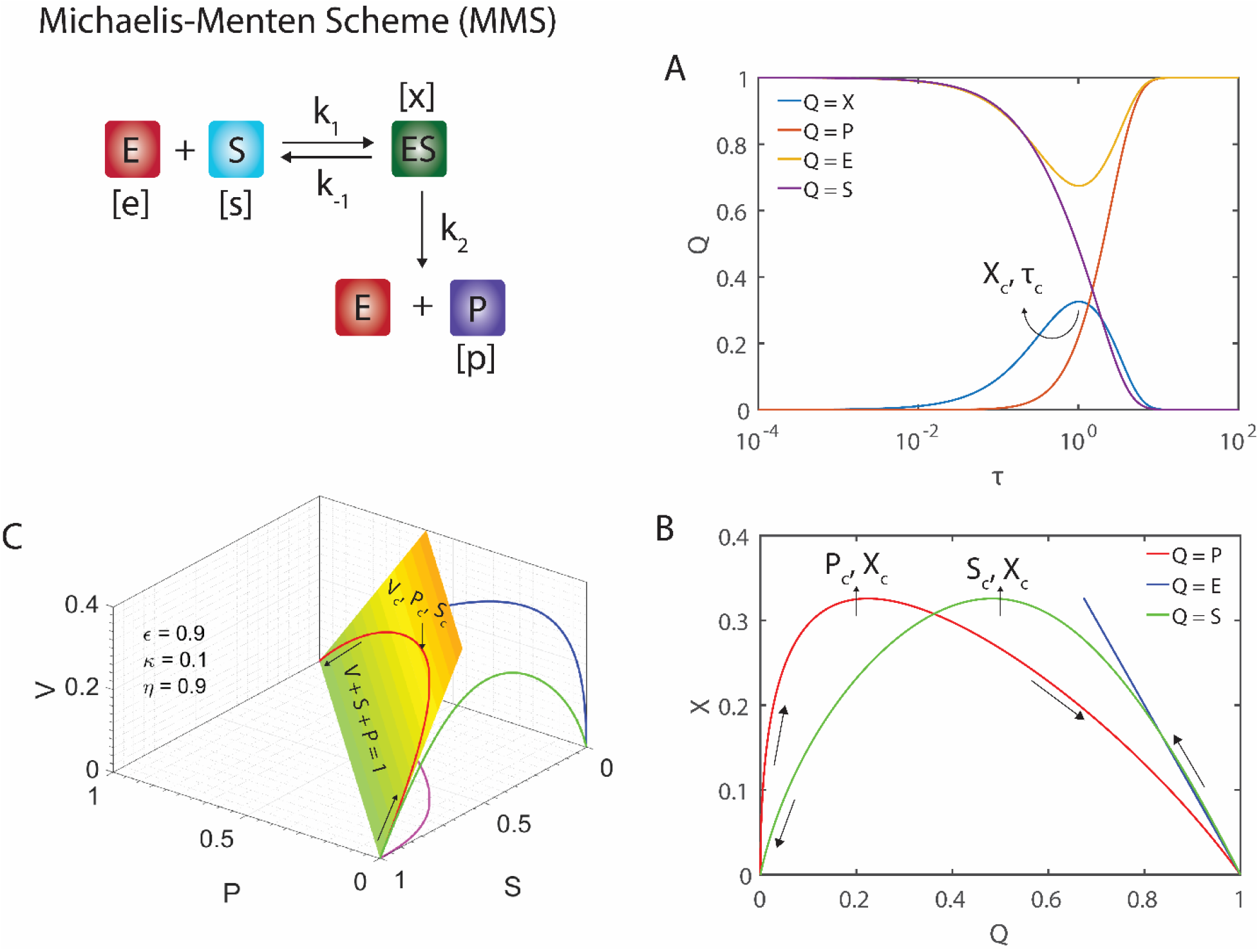
Michaelis-Menten enzyme kinetic scheme (MMS). Here the enzyme molecule first binds with the substrate to form enzyme-substrate complex in a reversible manner which irreversibly decays into free enzyme and product. Here [*e*], [*s*], [*x*] and [*p*] are the concentrations of enzyme, substrate, enzyme-substrate complex and product respectively. At *t* = 0, the initial conditions are (*e, s, p, x*) = (*e_0_, s_0_*, 0, 0). Apart from this we have conservation relations *e* = *e_0_* − *x* and *s* = *s*_0_ − *x* − *p*. **A**. Typical simulation trajectories of normalized concentrations (Eqs. 3) of enzyme (*E* = *e*/*e_0_*), substrate (*S* = *s*/*s*_0_), enzyme-substrate complex (*X* = *x*/*e_0_*) and product (*P* = *p*/*s*_0_) as a function of rescaled time with respect to *k_2_* i.e. *τ* = *k_2_t*. In the normalized space, the reaction velocity will be defined as *V* = *dP*/*dτ* = *εX*. Here the simulation settings for A, B and C are *ε* ~ 0.9, *η* ~ 0.9, *κ* ~ 0.1. Initial concentrations are set at (*E, S, X, P*) = (1, 1, 0, 0). Clearly there exists a steady state at time *τ_c_* where *dX*/*dτ* = 0. **B**. Dynamics of MMS in XP and XS spaces. At (*S_c_, P_c_*) we further have *dX*/*dS* = *dX*/*dP* = 0. **C**. Dynamics of MMS in VPS space. Red solid line is the parametric plot of the solution set (*V, P, S*) generated from the numerical solution of Eqs. 3 where green solid line is its projection on to VS plane, blue solid line is its projection on to VP plane and pink solid line is its projection over PS plane. All the trajectories of MMS in VPS space will lie on the plane defined by *V* + *S* + *P* = 1 which follows from the conservation relationships.

Several approximations were proposed in the light of experimental characterization of a single enzyme that is following MMS. The **standard quasi steady state approximation** (sQSSA) is widely used across several fields of biochemical research to characterize an enzyme. Analysis of MMS using sQSSA will yield two important parameters viz. *v*_max_ and *K_M_*. Here *v*_max_ = *k_2_e_0_* (Ms^−1^) is the maximum achievable reaction velocity (*v* = *dp*/*dt* = *k_2_x* where *p* is the concentration of product and *x* is the concentration of enzyme-substrate complex at time *t*) in MMS where *e_0_* is the initial enzyme concentration (M) and *K_M_* = (*k_−1_* + *k_2_*)/*k_1_* (M) is the Michaelis-Menten constant which characterizes the strength of binding of enzyme with its substrate. When the initial substrate concentration *s*_0_ is much higher than the initial enzyme concentration *e_0_* then sQSSA suggests an approximate expression for the reaction velocity as *v* = *v*_max_ *s* / (*K_M_*+ *s*) where *s* = (*s_0_ − x − p*) is the substrate concentration at time *t*. Under sQSSA conditions *x* will be much lesser than *s*_0_ and *p*. Under such conditions one can approximate the reaction velocity as *v* = *dp*/*dt* = −*ds*/*dt* which in turn yields the implicit form of integrated rate equation corresponding to sQSSA (Golicnik, 2013). Recently explicit expressions of integrated rate equation was obtained in terms of **Lambert’s W** functions (Golicnik, 2010; Golicnik, 2011a; Golicnik, 2011b; Schnell and Mendoza, 1997; Stroberg and Schnell, 2016). Apart from this a total QSSA (tQSSA) was also proposed where the approximation *s* = (*s_0_ − p*) was used to derive the steady state velocity of MMS. Using the experimental data on reaction velocity versus substrate concentration one finally obtains the MMS parameters *v*_max_ and *K_M_* via a general nonlinear least square fitting procedure. Several linearization techniques such as Lineweaver-Burk representation were also proposed (Atkins and Nimmo, 1975; Lineweaver and Burk, 1934). These in turn require a linear least square fitting procedure to obtain the MMS parameters.

Although it is easy to set the higher initial concentration of substrate than the enzyme under *in vitro* laboratory conditions, there are several situations such as single molecule enzyme kinetics (Grima et al., 2014) and other *in vivo* conditions where one cannot manipulate the ratio of substrate to enzyme concentrations much. Here the single molecule enzyme kinetics involves additional stochastic dynamics of enzyme, substrate and enzyme-substrate complex. Under *in vivo* conditions there are possibilities for the occurrences of comparable concentrations of enzyme and substrate or excess enzyme over substrate molecule. When the initial enzyme concentration is much higher than the initial substrate concentration, then a reverse QSSA (rQSSA) was proposed which predicted (Rami Tzafriri and Edelman, 2007; Tzafriri, 2003) an expression for the reaction velocity as *v* = *v_max_ s* / (*K_D_* + *s*) where *K_D_* = *k_−1_*/*k_1_*. Detailed studies suggested that successfulness of various QSSAs is strongly dependent on the timescale separation between the pre- and post-steady state dynamics of MMS (Borghans et al., 1996; Li et al., 2008). A short or transient pre-steady state timescale in which the enzyme-substrate complex *x* builds up from zero to steady state value quickly and a prolonged post-steady state timescales are ideal for applying various QSSA methods. Such conditions ensure accurate estimation of various MMS parameters (Borghans et al., 1996; Li et al., 2008).

Here steady state experiments will be set up by adding a series of known excess amounts of substrate to a known fixed amount of enzyme and then incubated at constant temperature and other environmental conditions for a fixed amount of time. Subsequently the enzyme reaction will be terminated by denaturing the enzyme. Increase in the product concentration over time with respect to increase in the initial substrate concentration will be recorded. When the timescale separation between pre- and post-steady state dynamics is high enough then sQSSA along with **reactant stationary assumption** (Stroberg and Schnell, 2016) will be applicable with *s* ~ *s*_0_ which yields the widely used sQSSA formula *v* ~ *v*_max_ *s*_0_ / (*K_M_* + *s*_0_). The reaction velocity *v* at a given initial substrate concentration *s*_0_ will be calculated by dividing the total product formed by the total experimental time. However as the enzyme catalyzes the reaction, concentration of substrate decreases and concentration of product increases monotonically along the time evolution. On the other hand, concentration of enzyme substrate-complex increases from zero to a maximum steady state value and then declines towards zero i.e. it will show up a typical turnover type pattern with time.

Since time is monotonically related to substrate as well as product concentrations, one can conclude that concentration of enzyme-substrate complex as well as reaction velocity will vary with both substrate and product concentrations in a turnover manner with a definite maximum. This means that *v* will be zero both at *s* = *s*_0_ (*p* = 0) as well as *s* = 0 (*p* = *s*_0_). In between these two boundaries there exists an optimum substrate concentration *s_c_* (or product concentration *p_c_*) at time *t* = *t_c_* at which *dv*/*dt* = 0 (and *s* = *s_c_* at which *dv*/*ds* = 0 or *p* = *p_c_* at which *dv*/*dp* = 0). This is precisely the steady state of MMS. Clearly sQSSA methods are applicable only for the range of substrate concentrations from *s* = *s_c_* to *s* = 0 (or *p* = *p_c_* to *p* = *s*_0_) which corresponds to the post-steady state dynamics of MMS. It is still not clear how the concentration of substrate as well as product influence the reaction velocity in the pre-steady state region of MMS.

In this article we first derive an ordinary perturbation expansion of the solution of MMS which is uniformly valid for both pre- and post-steady state regime of MMS. We identity an efficient combination of ordinary perturbation parameters which can give an accurate predictions about *s_c_, p_c_*, and various timescales like *t_c_* over a wide range of parameter space. We further derive an accurate expression for the dependency of *v* on s and *p* associated with the pre-steady state dynamics of MMS. In subsequent sections we reformulate the dynamical relationships among the variables (*v, s, p*) of MMS under normalized **vs, vp, ps** and **vps** phase spaces. We finally evaluate the applicability of various approximate expressions derived in this work along with the already existing ones from literature towards extracting various kinetics parameters from experimental data.

## 2. Theory

### 2.1. Scaling transformations of MMS and QSSAs

Let us denote the concentrations (mol/lit, M) of enzyme, substrate, enzyme-substrate complex and product associated with MMS shown in Fig. 1 as *e, s, x, p* respectively. Here we set the initial conditions at time *t* = 0 as (*e, s, x, p*) = (*e_0_, s_0_*, 0, 0) throughout this paper. Variables and parameters used throughout this paper are listed in Table 1. Noting the conservation relationships *e* = *e_0_* − *x* and *s* = *s*_0_ − *x* − *p*, the set of rate equations describing single substrate MMS can be written as follows.

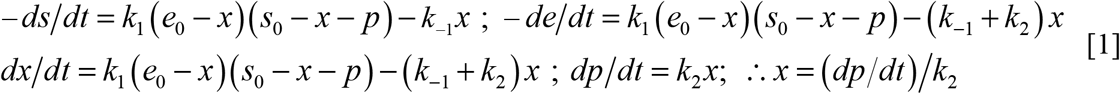

**Table 1:**
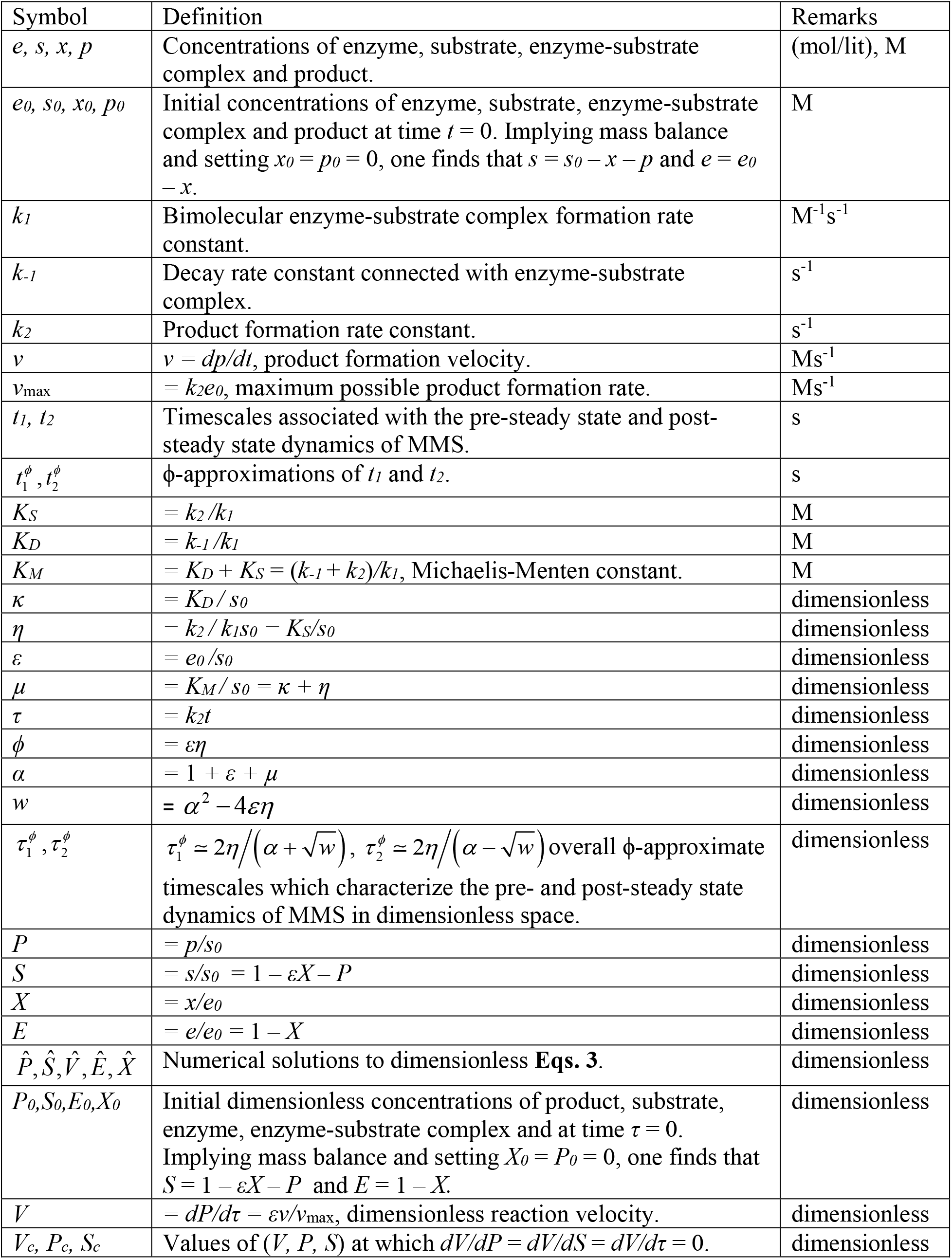
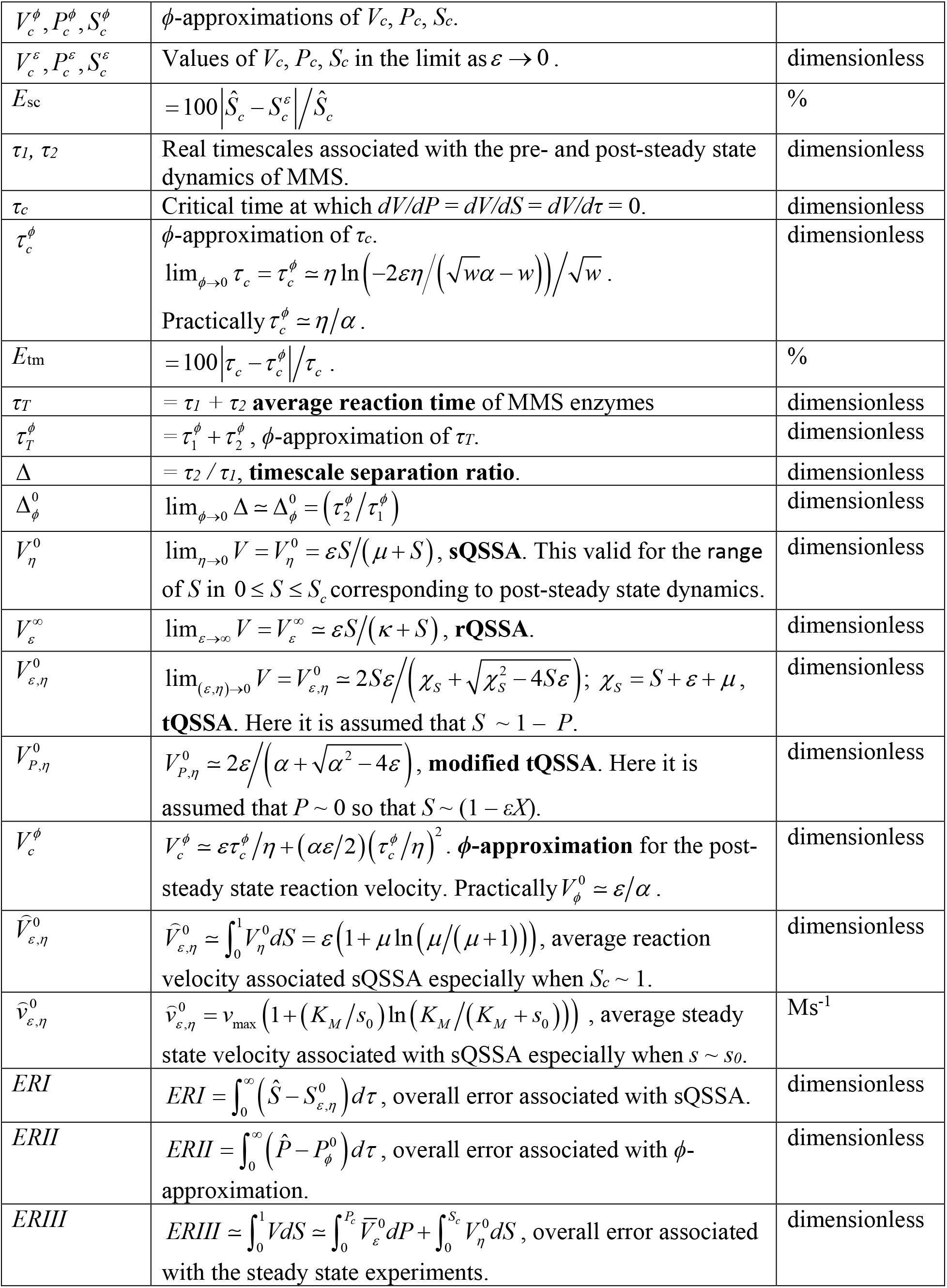
Various symbols and their definitions used in the theory section.

Analytic solution to the set of coupled nonlinear ordinary differential equations (ODEs) given in Eqs. 1 is unknown though one can expand the general solution in terms of ordinary (Murugan, 2002) or singular perturbation series (Dingee and Anton, 2008; Murray, 2002; Seshadri and Fritzsch, 1981). When [*dx*/*dt*] → 0 then one obtains the well-known **sQSSA** for the MMS relationship.

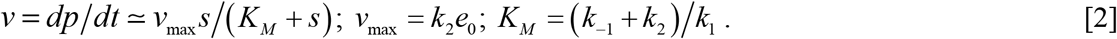

Here *v*_max_ (Ms^−1^) is the maximum possible reaction velocity and *K_M_* (M) is the Michaelis-Menten constant. We first perform the following rescaling of the dynamical variables to make the system of Eqs. 1 into a dimensionless one.

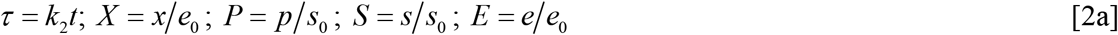

Using this scaling scheme, one can rewrite Eqs. 1 in a dimensionless form as follows.

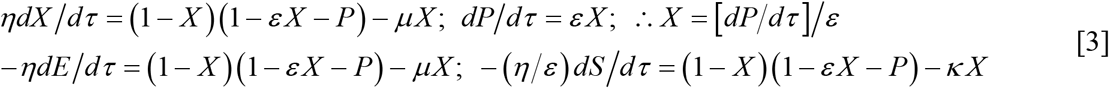

In this equation the dimensionless parameters (*ε, η, μ, κ*) are defined as follows.

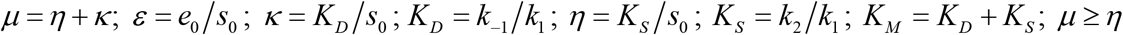

Here *μ* = *K_M_*/*s*_0_ and one finds the following mass balance and dynamical relationships.

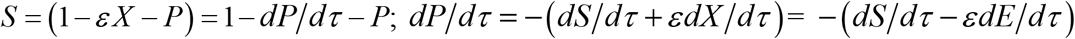

Numerically solved sample trajectories of Eqs. 3 for a given set of control parameters (*ε, κ, η*) are shown in Figs. 1A-C. Eqs. 3 undoubtedly suggest that the entire dynamics of MMS can be well characterized by the fundamental dimensionless parameters (*ε, η, κ*). Here *η* describes (**a**) how fast the kinetics of MMS approaches the steady state and (**b**) the extent of timescale separation between pre- and post-steady state dynamics. Parameter *κ* describes how fast the equilibrium associated with the formation of enzyme-substrate complex is achieved in the pre-steady state regime. Noting the fact that *V* = *dP*/*dτ* = *εX* is the reaction velocity of MMS, one finds that all the solution trajectories of Eqs. 3 will lie on the plane defined by *S* + *P* + *V* = 1 in VPS space (Fig. 1C). From Eqs. 1-3 one obtains the preliminary condition that is strictly required for the validity of sQSSA as *η* → 0 rather than [*dX*/*dτ*] → 0 as usually stated in the biochemical literature (Borghans et al., 1996). Actually one should note that the assumption [*dX*/*dτ*] = 0 is conceptually not a right one since [*dX*/*dτ*] ≠ 0 all through the dynamics of MMS enzymes except at the time point *τ* = *τ_c_*. Here one should note that *X* = 0 at *τ* = 0 as well as at *τ* → ∞. When *η* → 0 then one obtains 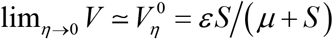 as sQSSA.

Under **reactant stationary assumption** (Stroberg and Schnell, 2016) one finds that *S* ~ 1 and subsequently we obtain the sQSSA formula 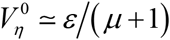 which is extensively used mainly to extract the values of *K_M_* and *v*_max_ from the steady state experimental data. On the other hand, the limiting condition *ε* → ∞ drives the term [*dS*/*dτ*] in the fourth equation of Eqs. 3 towards zero which finally results in the reverse QSSA (**rQSSA**) as 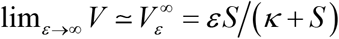. Here the dimensionless reaction velocity *V* of MMS can be transformed back in to the original velocity form *v* as *V* = *εv*/*v*_max_. From Eqs. 3 one finds the preliminary conditions required for the validity of the integrated rate equation associated with the sQSSA as follows.

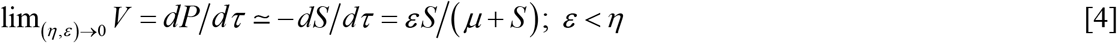

Since *S* = (1 − *εX* − *P*) the approximation used to derive Eqs. 4 i.e. *dP*/*dτ* ≃ −*dS*/*dτ* will be valid only when either *ε* or *X* tends towards zero. The main inconsistency in Eqs. 4 will arise especially when we set *η* → 0 which also drives [*dS*/*dτ*] towards zero. In this equation the required precise condition will be that *ε* → 0 much faster than *η* → 0. In other words, the condition *ε* < *η* is mandatory so that the term [*dS*/*dτ*] will not be driven towards zero while at the same time [*dX*/*dτ*] → 0.

The integral solution of the separable differential equation given in Eqs. 4 for the initial condition as *S* = 1 at *τ* = 0 can be explicitly written in terms of *Lambert’s W* function as follows (Golicnik, 2010; Rami Tzafriri and Edelman, 2007).

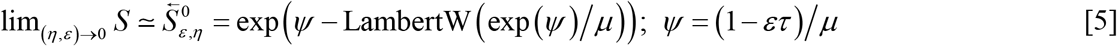

Here *y* = LambertW(*x*) is the solution of *ye^y^* = *x* (Corless et al., 1996). One can obtain this explicit solution (Eqs. 5 which is called as Schnell-Mendoza equation in the literature (Schnell and Mendoza, 1997)) by upon first solving Eq. 4 implicitly as (*S*/*μ*) exp (*S*/*μ*) = exp((1 − *ετ*)/*μ*)/*μ* and then inverting the left hand side of this equation in terms of Lambert’s W function. As in Figs. 2A and 2D, Eqs. 5 seems to fit the substrate depletion data very well especially in the post-steady state region of MMS (Golicnik, 2011b; Schnell, 2014). With the approximation given in Eqs. 4 and 5, one finds that lim_*ε*→0_ *P* = 1 − *S* and in general we have *P* ≤(1 − *S*). When (*ε, η*) → 0 then one finds the sQSSA value of *X* as lim_(*ε,η*)→0_ *X* ≃ *S*/(*μ* + *S*). Further one should note that Eqs. 5 will be valid only for the post-steady state dynamics of MMS (reverse arrow over *S* indicates this fact). We can define the error associated with Eqs. 5 in predicting the substrate depletion curve as **ERI** where 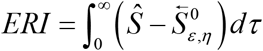. This is simply the total area associated with the gap in between the original and approximate trajectories. Figs. 3A-C show how **ERI** varies with respect to the parameters (*ε, κ, η*). Here *Ŝ* is the numerical solution to Eqs. 3 for a given set of parameters (*ε, η, κ*) with the initial condition as *S* = 1 at *τ* = 0. In the later sections we will show that one can derive expression for the product evolution curve (*P*) similar to Eqs. 5 without making assumption such as *P* ≃ 1 − *S* etc.

**FIGURE 2:**
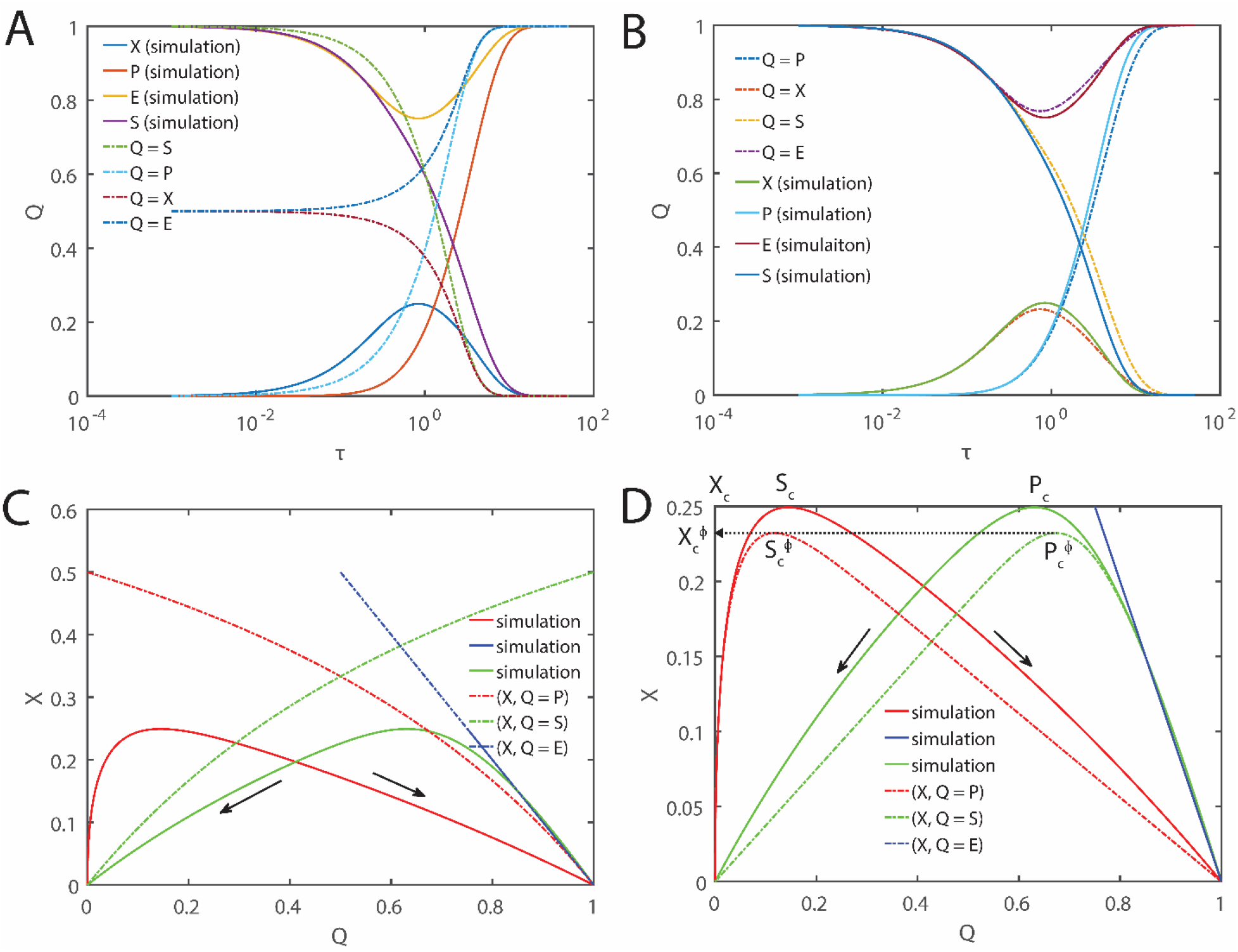
Comparison of integrated form of the rate equation associated with sQSSA (**A** and **C**) and *φ*-approximation (**B** and **D**) with the numerical simulation results. Solid lines are the numerical integration of Eqs. 3 using Euler scheme with Δτ ~ 10^−5^ for a total time of *τ* ~ 100. Dotted lines are the respective approximations. For integrate rate equation associated with sQSSA we used Eq. 5 and for *φ*-approximation we used Eq. 8. Here the initial settings at *τ* = 0 are *S_0_* = 1, *P_0_* = 0, *X_0_* = 0, *E_0_* = 1. Further we have set the parameters *ε* ~ 0.9, *κ* ~ 0.1 and *η* ~ 0.9. **C** and **D** describes the dynamics in (*X*, [*P, S, E*]) spaces. Clearly integrated rate equation works well only in the post-steady state region of MMS. On the other hand *φ*-approximation works well only in the pre-steady state region of MMS. Here (*X_c_, P_c_, S_c_*) are the steady state original values of *X, P* and *S* respectively and 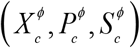 are the corresponding *φ*-approximations.

**FIGURE 3:**
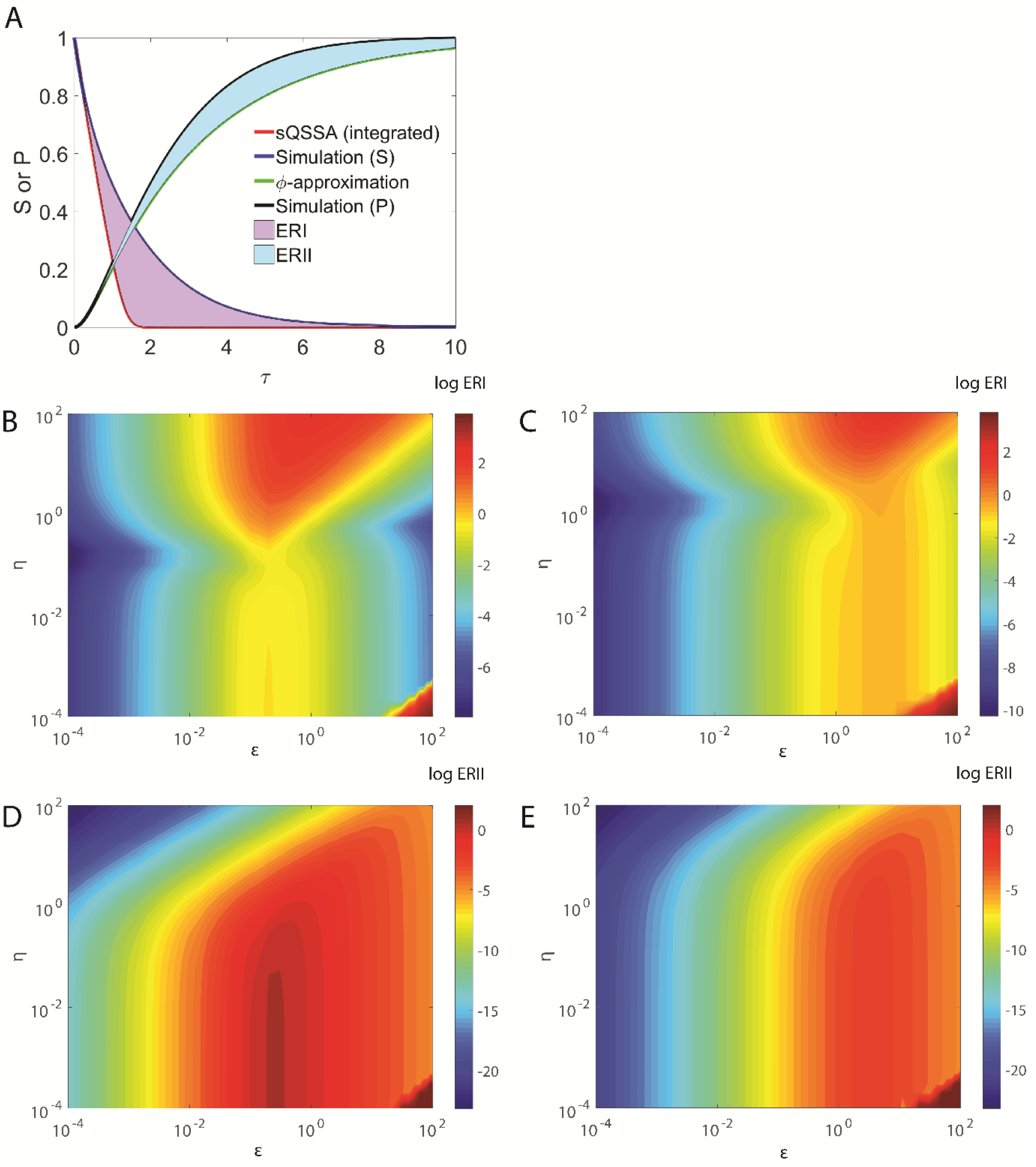
Errors associated with the integrated form of sQSSA where the approximation *dP*/*dτ* ≃ −*dS*/*dτ* is used (**ERI**) i.e. *P* ~ 1 − *S* (this will be true only when *ε* = 0 or *X* = 0 since originally we have *P* = 1 − *εX*− *S*) and *φ*-approximation (**ERII**) with respect to changes in the parameters (*ε, η, κ*). **A**. Here the settings are *ε* ~ 0.9, *η* ~ 0.9, *κ* ~ 0.1. Here errors ERI and ERII are the areas confined between the approximations and original simulation trajectories. The solid red line is the integrated sQSSA approximation given by Eqs. 5 with the initial condition as *S* = 1 at *τ* = 0 and green solid line is the φ-approximation given by Eq. 8 with the initial condition as *P* = 0 at *τ* = 0. **B.** ERI where *κ* ~ 0.1. **C**. ERI were *κ* ~ 1. **D**. ERII where *κ* ~ 0.1. **E**. ERII where *κ* ~ 1.

Although Eqs. 5 approximates the substrate depletion curve nicely, it fails to model the reaction velocity data (*V, τ*) especially in the pre-steady state region of MMS. Noting the fact that the reaction velocity is approximated here as *V* = *dP*/*dτ* ≃ −*dS*/*dτ* while deriving Eqs.5, when *τ* < *τ_c_* then one finds that 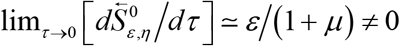 which is contradictory to the fact that we have *V* = 0 at *τ* = 0 (because *X* = 0 at *τ* = 0). This discrepancy mainly comes from the fact that [*dV*/*dτ*] ≠ 0 except at initial, end and steady state of MMS dynamics. On the other hand Eqs. 5 assume [*dV*/*dτ*] = 0 throughout the dynamics of MMS and there is no way of setting the initial condition for *V* in such case since it is a first order ODE. Eqs. 5 can also be written in a simple form as 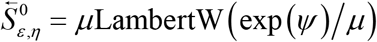 which follows from the property of the **Lambert’s W** function as LambertW(*u*) = *u* exp(−LambertW(*u*)).

### 2.2. Solution to MMS via ordinary perturbation series and φ-approximations

Several ordinary and singular perturbation expansions (Dingee and Anton, 2008; Murugan, 2002; Schnell and Mendoza, 1997; Seshadri and Fritzsch, 1981) and expansions based on slow manifold theories (Fraser, 2004; Roussel and Fraser, 2001; Roussel and Tang, 2006) were proposed for the general solution of Eqs. 1. One can generalize these ideas based on how the dimensionless parameters (*ε, η, κ*) approach towards zero. We first consider the following scaling transformation.

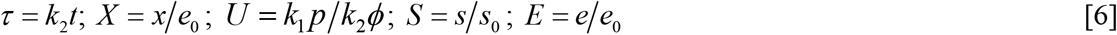

In this current work we propose the product *φ* = *εη* as the ordinary perturbation parameter rather than individual (*ε, η*) as in case of various QSSAs. This means that the zeroth order perturbation term obtained with this scaling scheme will be superior to other methods which use either *ε* → 0, *η* → 0 as in sQSSA or both (as in Eqs. 5) in an independent manner. With this transformation and upon defining *α* = 1 + *ε* + *μ*, Eqs. 1 can be rewritten as a second order nonlinear ODE as follows.

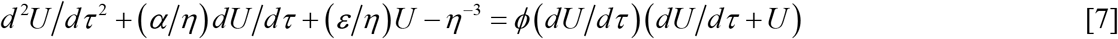

Here *φ* is the ordinary perturbation parameter which multiplies the nonlinear terms present on the right hand side of Eq. 7 and rescaled variable *U* is connected with *P* as in Eqs. 6 via the scaling transformation rule *P* = *εη*^2^*U*. Eqs. 3 have been expressed as second order differential equation earlier (Eq. 11 in Ref. (Morales and Goldman, 1955)). A systematic ordinary perturbation scheme corresponding to Eqs. 3 was first formulated by Murugan in Ref. (Murugan, 2002) and we call Eq. 7 and related transformations as Murugan equations. Eq. 7 and associated solutions are the central results of this paper from which we will derive several types of approximations by setting different limiting conditions for the control parameters. Firstly we denote the expansion of the solution of MMS over the parameter *φ* around *φ* = 0 as ***φ*-approximation**. Following the standard perturbation expansion of *U* in terms of undetermined gauge functions (Murdock, 1991; Murugan, 2002) as *U* = *U*_0_ + *U*_1_*φ* + *U*_2_*φ*^2^ + …, the zeroth order perturbation term *U*_0_ associated with the limit as *φ* → 0 can be obtained in terms of *P* of Eqs. 3 as follows.

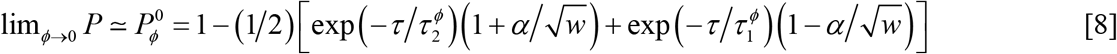

In this equation the exponential terms and other parameters are defined as follows.

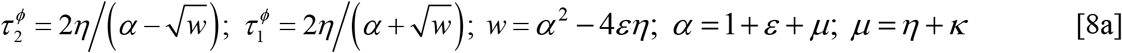

Here 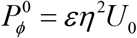. Using Eq. 8 one can further derive the expressions for *X* and *S* in the *φ*-approximation limit as 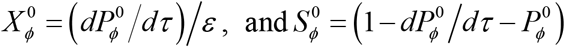. We can define the error associated with Eq. 8 in predicting the product generation or progress curve as **ERII** where 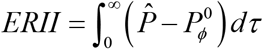. Here 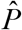 is the numerical solution to Eqs. 3 for a given set of parameters (*ε, η, κ*) with the initial condition as *P* = 0 at *τ* = 0. Since there is a *η*^−3^ term in the differential equation corresponding to 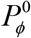 in the limit as *φ* → 0 i.e. left hand side of Eq. 7, Eq.8 will be accurate enough only in the pre-steady state region of MMS which only requires the condition as *ε* → 0. These results are clearly shown in Figs. 2B and D. Further Figs. 3A, D and E show how **ERII** varies with respect to the parameters (*ε, κ, η*).

#### 2.2.1. Steady state timescale

In Eq. 8 the exponential term 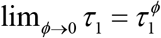 decides how fast the dynamics of MMS enzymes approaches the steady state starting from time *τ* = 0 and the term 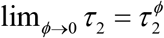 decides how fast the final enzyme-substrate complex decays into free enzyme and product. This is evident from the fact that 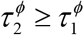. Since *dP*/*dτ* = *εX* one can show that [*dX*/*dτ*] will be zero at some critical time *τ_c_* which can be written explicitly in the limit *φ* → 0 as follows.

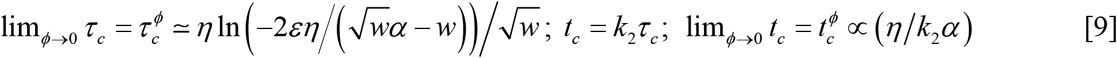

Here (*w, α*) are defined as in Eqs. 8a. Explicit expressions for the timescales *τ*_1_, *τ*_2_ and *τ_c_* are not known. From Eq. 8 one can conclude that when *τ* = *τ_c_* then *X* will attain a maximum where the reaction velocity *V* also becomes a maximum so that at time *τ_c_* we have *dV*/*dτ* = *dX*/*dτ* = 0. The steady state of MMS dynamics occurs exactly at this time point. The percentage error associated with the φ-approximation of *τ_c_* by 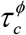 can be defined as 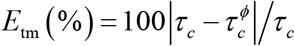. Figs. 4A-D show how *E*_tm_ varies with respect to changes in the control parameters (*η, ε, κ*). Here the true value of *τ_c_* can be obtained by numerically solving Eqs. 3 for a given set of parameters (*ε, κ, η*) with the initial conditions as (*S, P, X, E*) = (1, 0, 0, 1) at *τ* = 0 and then finding the time point at which [*dX*/*dτ*] = 0 by numerical differentiation of the trajectory of *X*. Here *τ*_1_ and *τ*_2_ are the actual timescales associated with the pre- and post-steady state dynamics of MMS enzymes (in the dimensionless space) and, 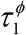 and 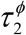 are their corresponding *φ*-approximations.

**FIGURE 4:**
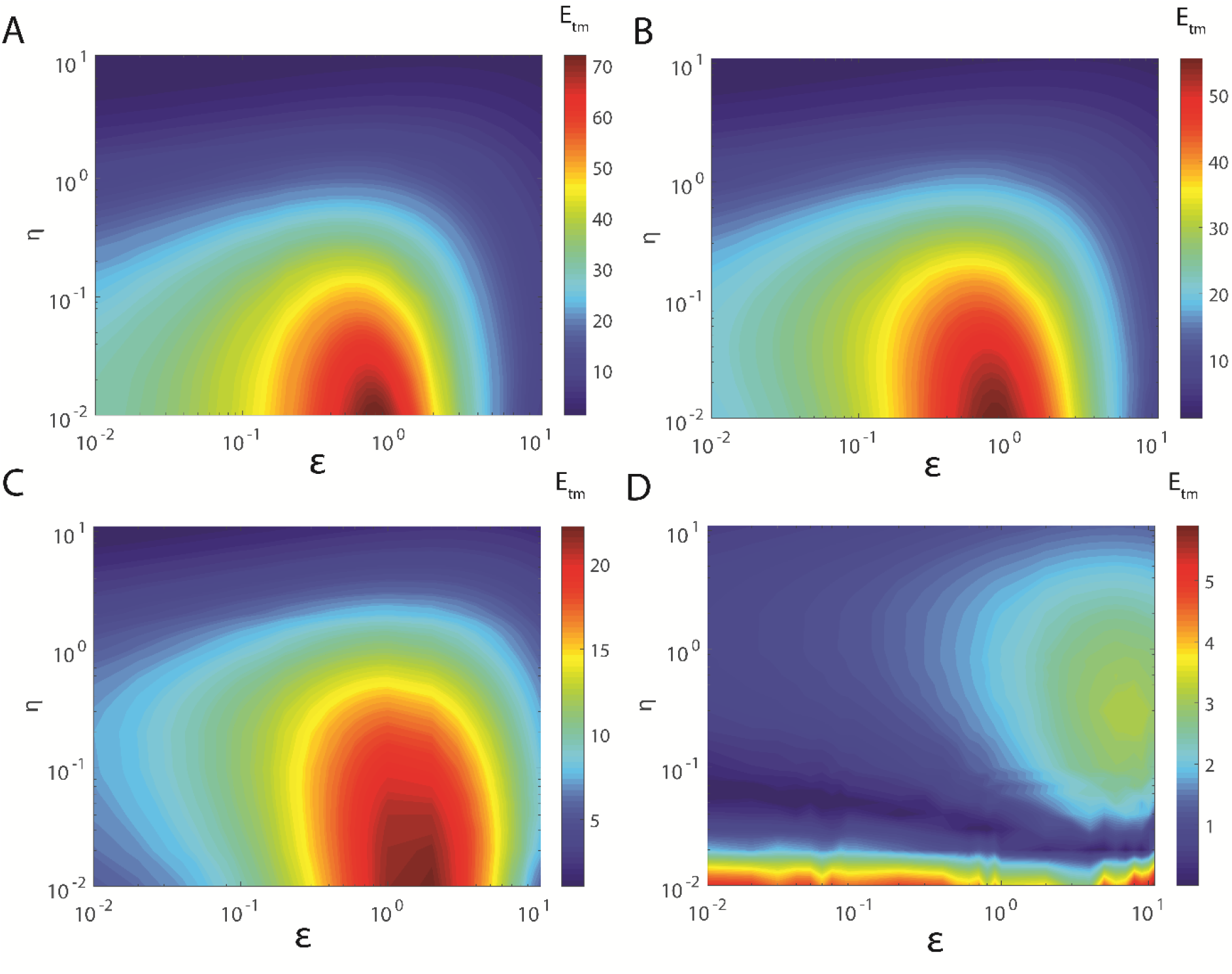
Error (*E*_tm_) associated with the *φ*-approximation i.e. 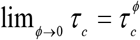 of the time that is required by MMS enzymes to attain the steady state. Explicitly this percentage error can be written as 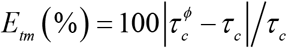 where *τ_c_* is the original value obtained from numerical simulation of Eqs. 3 and 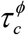 is its φ-approximation obtained from Eqs. 9. To obtain the original value of *τ_c_* one needs to integrate Eqs. 3 numerically for a given set of parameters (*κ, η, ε*) with the initial conditions (*X, P, S*) = (0, 0, 1) at *τ* = 0 and then identify the time point at which [*dX*/*dτ*] = 0 by numerical differentiation. At this time *τ* = *τ_c_*. **A**. *κ* = 0.01. **B**. *κ* = 0.1. **C**. *κ* = 1. **D**. *κ* = 10.

#### 2.2.2. Timescale separation ratio

Additionally one finds that *τ*_1_ ≤ *τ_c_* ≤ *τ*_2_ and 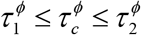. Using these timescale quantities one can define an effective **timescale separation ratio** that is available for conducting steady state *in vitro* experiments as 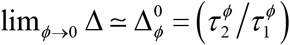. Figs. 5A-D show how 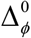 varies with respect to changes in the parameters (*η, ε, κ*). Explicitly one can write down the expression for this ratio as 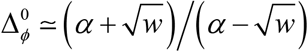. This clearly suggests that the effective timescale separation ratio can be well manipulated by the parameters (*ε, η, κ*). Particularly we have the limiting conditions as 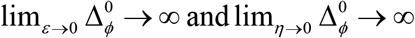. Since we also have the limits as 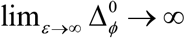 and 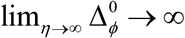, there should be an optimum point with respect to variation in both (*ε, η*).Upon solving 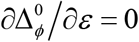 for *η* one finds that there exists a minimum of 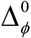 at *ε*_m_ = 1 + *η* + *κ*. Similarly upon solving 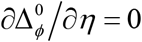 for *η* one finds that there exists a minimum of 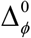 at *η*_m_ = 1 + *ε* + *κ*. In the later sections of this paper, we will show that the overall error associated with the sQSSA methods in predicting the reaction velocity of MMS will be a maximum at these optimum points *ε*_m_ and *η*_m_. That is to say, the error level of sQSSA methods in predicting the steady state velocity of MMS enzymes increases along with the decreasing timescale separation ratio. Here one should also note the limit with respect to *κ* as 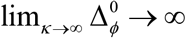. The explicit expressions for the minimum value of 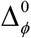 i.e. 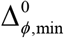 can be written in terms of *ε*_m_ and *η*_m_ as follows.

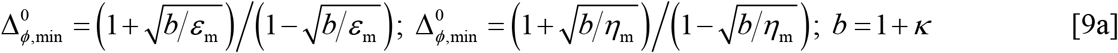

**FIGURE 5:**
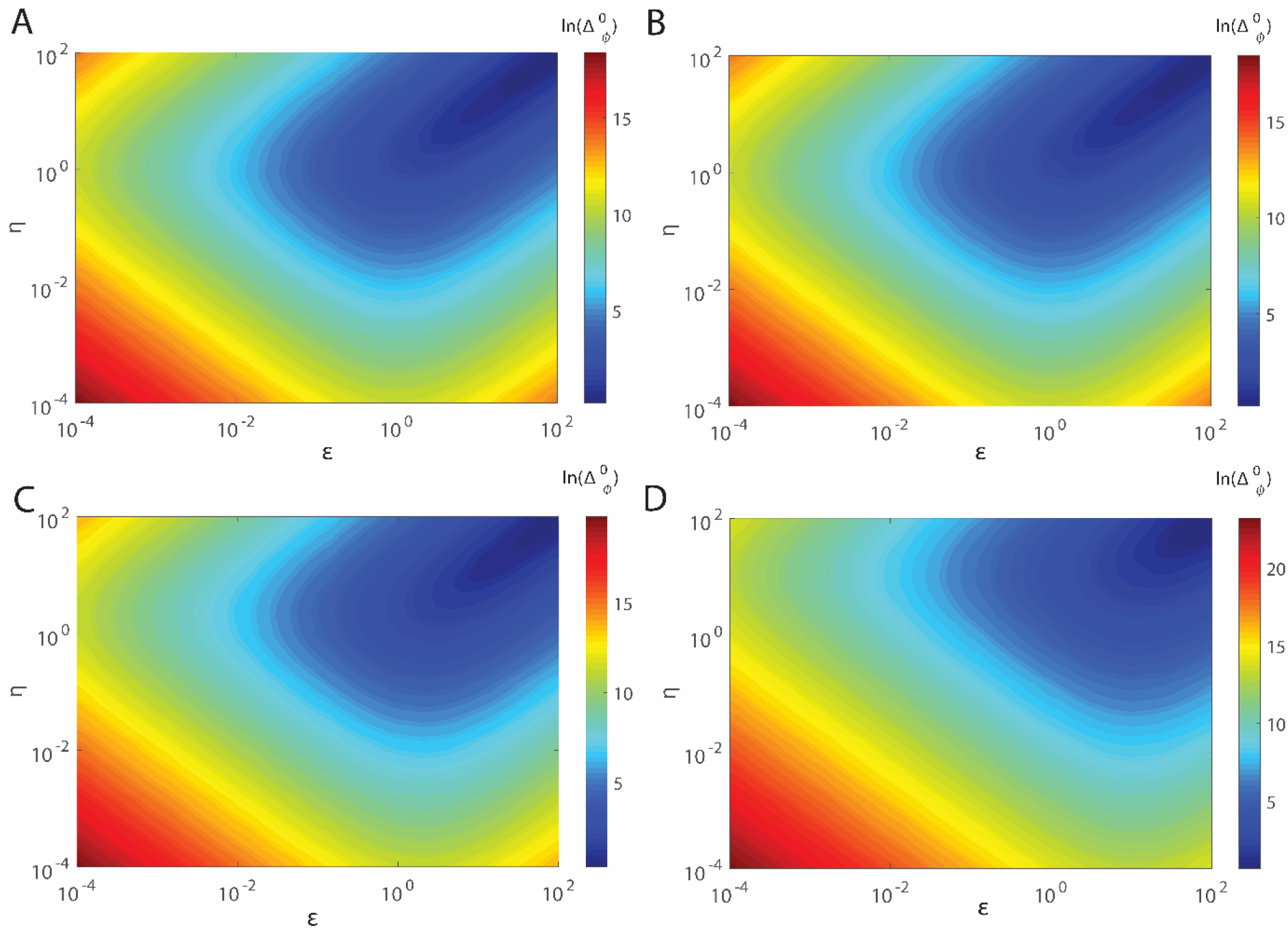
Effective timescale separation ratio 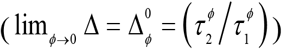 associated with MMS enzymes as predicted by the *φ*-approximation (Eqs. 8). Successfulness of steady state experiments in extracting various Michaelis-Menten parameters such as *K_M_* and *v*_max_ seems to be directly proportional to the extent of this timescale separation ratio. Dynamical efficiency of MMS enzyme in clearing toxic substrates from the biological medium will be inversely proportional to this timescale separation ratio. Clearly 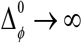 as *ε* → 0, *η* → 0 or *φ* → 0. **A**. *κ* = 0.01. **B**. *κ* = 0.1. **C**. *κ* = 1. **D**. *κ* = 10.

For sufficiently small values of *φ* one finds from Eq. 9 that *t_c_* ≃ 1/*k*_1_(*s*_0_ + *e*_0_ + *K_M_*). Borghans et.al. in Ref. (Borghans et al., 1996) have derived similar expression for the steady state timescale *t_c_* using QSSA arguments. Expressions for these pre- and post-steady state timescales in terms of original MMS variables can be written as follows.

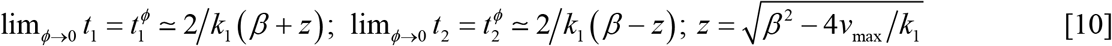

Here *β* = (*s*_0_ + *e*_0_ + *K_M_*). One should note that *t*_1_ and *t*_2_ are the actual timescales associated with the pre- and post-steady state dynamics of MMS (in terms of original variables) and, 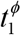 and 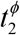 are their corresponding *φ*-approximations. Hence obtained values of these timescales are close to the earlier suggested values as (*t*_1_, *t*_2_) ≃ (1/*k*_1_ (*s*_0_ + *K_M_*),(*s*_0_ + *K_M_*)/*v*_max_) which were actually obtained using tQSSA arguments (Borghans et al., 1996). When *ε* → 0 then these timescales will transform as (*t*_1_,*t*_2_) ≃ {1/2*βk*_1_, ∞}. This follows from the fact that we have 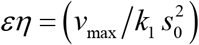 in Eqs. 10. These results suggest that one can increase the steady state *in vitro* experimental range of a single substrate MMS enzyme mainly by decreasing *ε* and *μ* which can be achieved mainly by raising *s*_0_.

#### 2.2.3. Average reaction time and catalytic efficiency

The kinetic efficiency of a MMS enzyme can also be measured by the **average reaction time** that is required to completely convert the initial amount of substrate into the respective product i.e. *τ_T_* = *τ*_1_ + *τ*_2_. Although exact expression for *τ_T_* is not known for MMS enzymes one can use the corresponding *φ*-approximation as 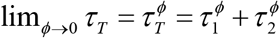. Explicitly one obtains the *φ*-approximation using Eqs. 8 as 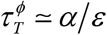 from which one finds following limiting conditions with respect to *ε* as 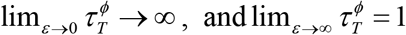. Similarly one finds the limiting conditions with respect to *η* as 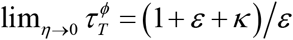, and 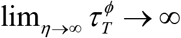. In terms of original MMS variables we have 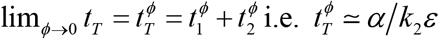. The average time required by a single enzyme molecule to convert a single substrate in to product 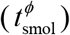 can be calculated from 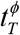 as 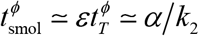 from which one can straightforwardly derive the single molecule Michaelis-Menten equation as 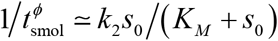 (Kou et al., 2005).

Here one should note that the total timescale quantity (*t_T_*) is a critical parameter especially for the scavenging enzymes like Cytochrome P450s (though this is not a single substrate MMS enzyme) over toxic substrates (Ortiz De Montellano, 2016). Actually the overall efficiency of the detoxifying enzymes will be inversely proportional to *t_T_*. This is because those toxic substances released inside the cellular cytoplasm during various metabolic activities need to be immediately captured and subsequently removed or destroyed before they cause damage to other cellular organelles. One can define the **catalytic efficiency** of a MMS enzyme as inverse of the total reaction time as 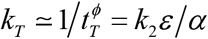 and one finds the limiting conditions as lim_*ε*→∞_ *k_T_* → *k*_2_, lim_*s*_0_→∞_ *k_T_* → 0 and lim_*K_M_*→∞_ *k_T_* → 0. In terms of original variables we have *k_T_* ≃ *k_E_e*_0_. Here we have defined *k_E_* = *k*_2_/(*s*_0_ + *e*_0_ + *K_M_*) as the measure of catalytic efficiency of a single substrate MMS enzyme which has the dimension of M^−1^s^−1^. It is interesting to note that this is similar to the expression *k_E_* = *k_cat_*/*K_M_* (where *k*_cat_ = *k_2_* in the present context) which is widely used in the literature especially for measuring and comparing the catalytic efficiencies of the single substrate MMS enzymes (Alberts, 2002; Stryer, 1988; Voet and Voet, 1995). Our results suggest that the widely used definition for the catalytic efficiency i.e. *k_E_* = *k*_cat_/*K_M_* will be valid only under dilute solution conditions where *K_M_* ≫ (*s*_0_, *e*_0_).

#### 2.2.4. Steady-state reaction velocity

The most appropriate way of performing the steady state experiments will be via setting the parameter *φ* → 0 rather than by setting *ε* → 0 and *η* → 0. This condition is close to the earlier proposed overall condition for the validity of sQSSA (Borghans et al., 1996). In terms of the dimensionless variables of this current work, the condition proposed for the validity of sQSSA in Ref. (Borghans et al., 1996) can be rewritten as (*φ*/(1 + *μ*)^2^) ≪ 1 which will be obviously true in the limit as *φ* → 0. The steady state reaction velocity of MMS can be obtained by substituting 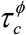 of Eqs. 9 into 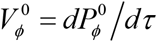 where 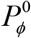 is defined as in Eq. 8. Explicitly one can write the expression for the steady state reaction velocity corresponding to the *φ*-approximation as follows.

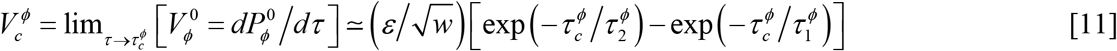

Here 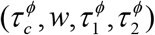 are defined as in Eqs. 8 and 9. In the limit as *φ* → 0 Eq. 11 will be more accurate in predicting the steady state reaction velocity than the integrated rate equation associated with sQSSA. Upon expanding the right hand side of 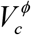 in Eq. 11 in a Macularin series with respect to 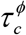 and terminating the series beyond the second order term we obtain 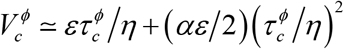 which accurately represents the steady state reaction velocity of MMS especially when the pre-steady state timescale is very small. For small values of *ε* one finds that 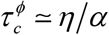 from which one obtains 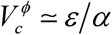. In terms of the original variables we find that 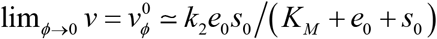.

It is interesting to note that we have obtained this expression for the reaction velocity without imposing the **reactant stationary assumption** over sQSSA i.e. *s* ≃ *s*_0_ *S* ~ 1. Further, this reaction velocity expression will behave as 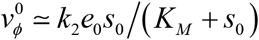 particularly when *e*_0_ ≪ *s*_0_. It will behave as 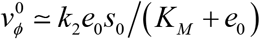 when *e*_0_ ≫ *s*_0_. These results are in fact consistent with the recent studies of Kargi in Ref. (Kargi, 2009) and Bajzer and Strehler in Ref. (Bajzer and Strehler, 2012). One also should note that unlike Eqs. 5, the *φ*-approximation given by Eq. 8 will be valid for both pre- and post-steady state region of MMS. Similar to Eq. 11 one can substitute 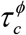 into the expression for 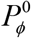 as in Eq. 8 to obtain the approximate steady state product levels 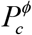 and subsequently one finds 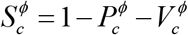 for the approximate steady state substrate levels.

### 2.3. Dynamics of MMS in XQ spaces where Q = (P, E, S)

Following the scaling transformation as in Eqs. 2a and noting the definition of dimensionless reaction velocity *V* = *dP*/*dτ* = *εX*, Eqs. 3 and 7 can be rewritten in XQ spaces where Q = P, E, S as follows (Figs. 6).

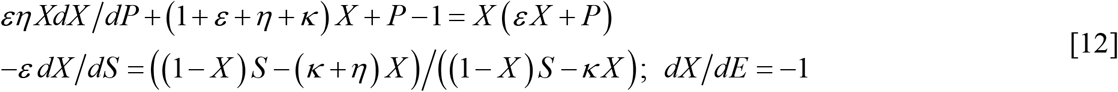

**FIGURE 6:**
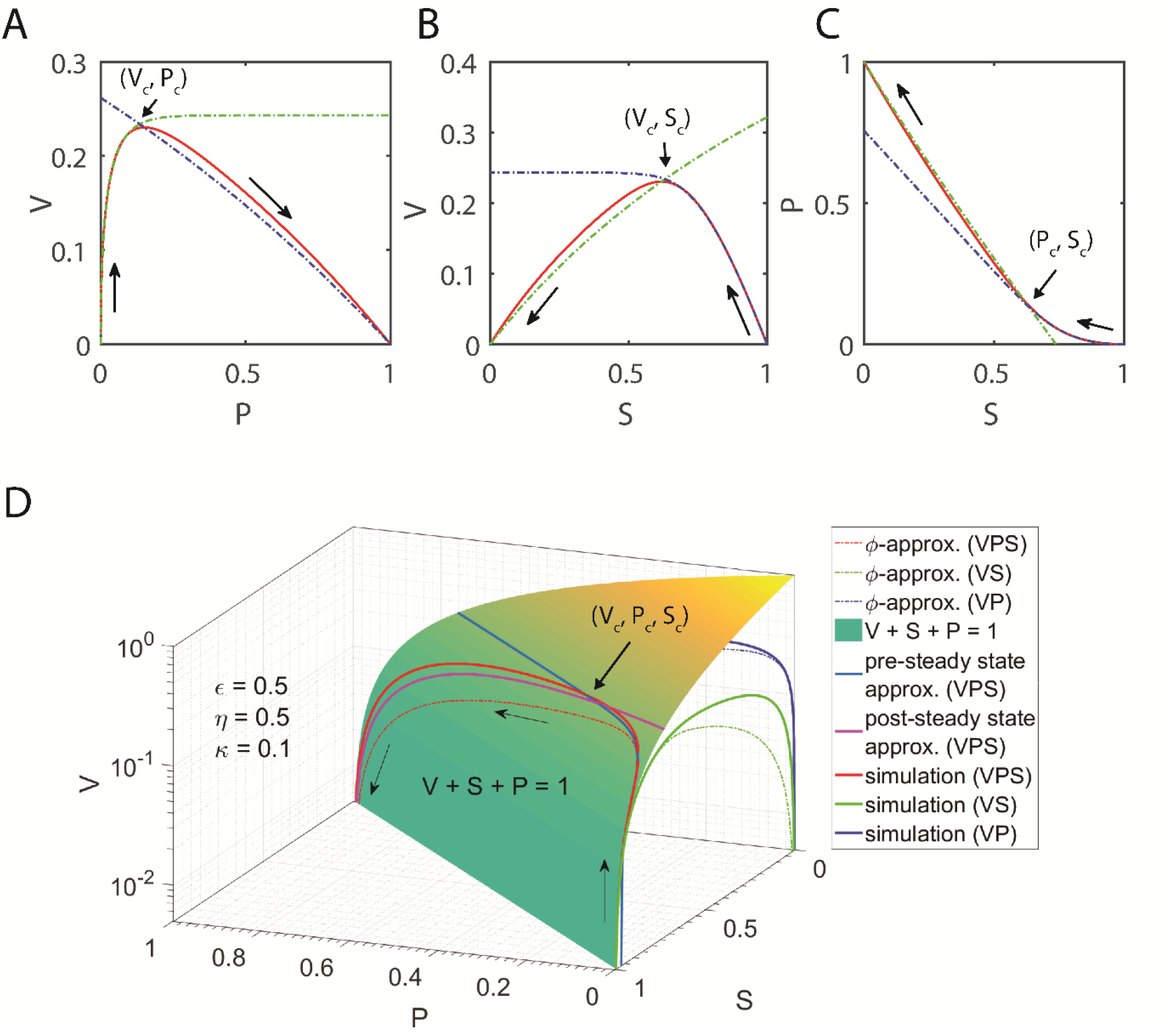
Dynamics of MMS in VP, VS, PS and VPS spaces. Settings for **A, B** and **C** are *ε* = 0.9, *η* = 0.9, *κ* = 0.9. **A**. Red solid line is the numerical integration of Eq. 24, dotted green line is the pre-steady state approximation 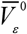 as in Eq. 26 and dotted blue line is the post-steady state approximation 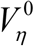 as in Eq. 16. Clearly these two approximations intersect at the steady state critical point (*V_c_, P_c_*). **B**. Red solid line is the numerical integration of Eq. 19, dotted blue line is the pre-steady state approximation 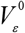 as in Eq. 21 and dotted green line is the post-steady state approximation 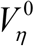 as in **sQSSA**. Clearly these two approximations intersect at the steady state critical point (*V_c_, S_c_*). **C**. Red solid line is the numerical integration of Eq. 30, dotted green line is the post-steady state approximation 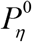 (as defined below Eq. 30) and dotted blue line is the pre-steady state approximation 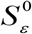 as in Eq. 33. Clearly these two approximations intersect at the steady state critical point (*S_c_, P_c_*) in SP space. **D**. Data on the plane surface defined by *V* + *P* + *S* = 1 (*V* is in log-scale for better visibility). Red solid line is the numerical integration of Eq. 35 with the initial condition *V* = 0 at *S* = 1 and *P* = 0. Dotted red line is the parametric plot made with *φ*-approximation. Rose solid line is the parametric representation 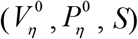 corresponding to the post-steady state approximation as in Eq. 38 and solid navy line is the parametric representation 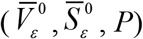 for the pre-steady state approximation as in Eq. 37. **VS plane**. Green solid line is the numerical integration of Eq. 19 and dotted green line is the parametric plot made with *φ*-approximation. **VP plane.** Blue solid line is the numerical integration of Eq. 30 and dotted blue line is the parametric plot made with *φ*-approximation.

From the first equation of Eqs. 12, in the limit as *ε* → 0 one can obtain *X* as *X* ≃ *ξ*/(*μ* + *ξ*) where *ξ* = 1 − *P*. Noting that *X* = [*dP*/*dτ*]/*ε*, one can derive a separable type differential equation which is valid in the *ε* → 0 as [(*μ* + 1 − *P*)/(1 − *P*)] *dP* ≃ *εdτ*. The integral solution of this nonlinear ODE will be similar to that of the integrated form of sQSSA given in Eqs. 5. This can be written explicitly as follows.

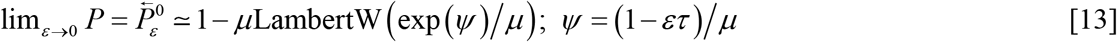

Upon comparing this equation with Eqs. 5 (which strictly requires the conditions that *ε* → 0 and *η* → 0) one finds that 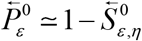 and clearly Eqs. 13 (which requires only the condition that *ε* → 0) will be valid only for the post-steady state region of MMS. When we have *η* → 0 then using the first equation in Eqs. 12 one can derive an approximate expression for *X* in terms of *P* as follows.

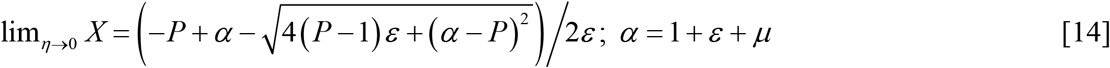

Upon solving the quadratic equation *αX* + *P* − 1 = *X* (*εX* + *P*) for *X* one needs to impose the condition that *X* ≤ 1 and select the appropriate root accordingly. This type of total QSSA (**tQSSA**) will be generally derived by substituting *S* = 1 − *P* (which in turn requires the condition that *ε* → 0 or *X* → 0) in the first equation of Eqs. 3 and subsequently solving it for *X* in terms of *S* in the limit as *η* → 0. As a result one obtains the following expression for the reaction velocity corresponding to tQSSA (Borghans et al., 1996; Tzafriri, 2003).

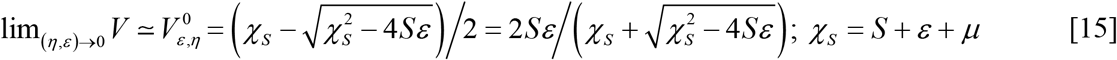

One should note that Eqs. 15 works very well in both the limits as (*ε, η*) → 0 independently, and also more accurate than the sQSSA one i.e. *V* = *εS*/(*μ* + *S*) where *S* = 1 − *P* in XP or VP space that was obtained from Eqs. 12 under the condition that *ε* → 0. Expression similar to tQSSA as given in Eqs. 15 has been derived recently using the approximation as *S* ≃ (1 − *εX*) by ignoring *P* in the first equation of Eqs. 3 and subsequently solving for the steady state of *X* by setting *η* → 0. The steady state velocity can be expressed as 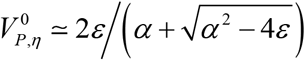. Borghans et.al in Ref. (Borghans et al., 1996) and Tazfriri in Ref. (Tzafriri, 2003) (e.g. Eq. 11 in Ref. (Tzafriri, 2003)) used similar type of assumption of ignoring *P* term from the definition of *S* while formulating their tQSSA. This is somewhat a partial reactant stationary approximation. Clearly 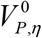 will be a function of *K_M_, e_0_* and *s*_0_. Bajzer and Strehler (Bajzer and Strehler, 2012) have shown that 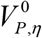 (Bajzer and Strehler, 2012) predicts the original steady state reaction velocity very well with minimum amount error under both the conditions as *ε* → 0, *ε* → →. In this context one can derive an expression for the reaction velocity *V* in terms of *P* using Eqs. 14 as follows.

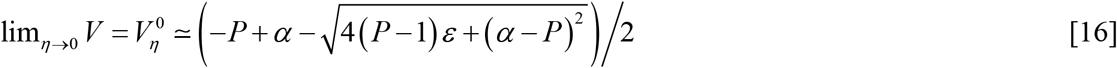

Upon expanding the right hand side terms in Eq. 16 in a Macularin series around *ε* = 0 and noting that *S* = 1 − *P* under such conditions, one can find the following expansion for the reaction velocity equation associated with the sQSSA of MMS.

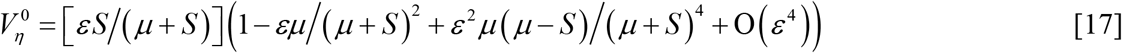

Expression similar to Eq. 17 has been derived earlier by Schnell and Maini in Refs. (Schnell and Maini, 2000; Schnell and Maini, 2002). Upon ignoring the higher order *ε* terms in the limit as *ε* → 0 one recovers 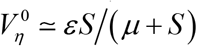. These results suggest that tQSSA underestimates the reaction velocity of MMS especially for large values of *ε*. Noting that *P* + *S* + *V* = 1, one can derive an expression from Eq. 16 which connects *S* and *P* in the limit *η* → 0 as follows.

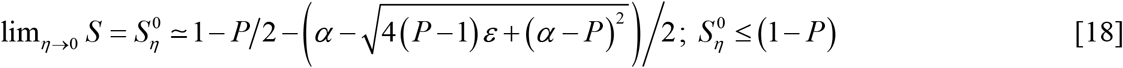

This equation will be valid only for the post-steady state regime of MMS which is evident from the fact that at *P* = 1 we have *S* = 0. However upon insisting the initial condition for pre-steady state regime as *P* = 0 one finds that *S* ≠ 1.

### 2.4. Dynamics of MMS in the VS space

Noting that *V* = *εX* from Eqs. 12 one can derive the following dynamical equation of MMS in the VS space (Fig. 6B).

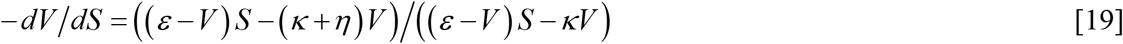

This equation suggests that there exists an optimum set of points (*V_c_, S_c_*) in the VS space at which [*dV*/*dS*] = 0. This will be exactly the steady-state in (*S, τ*), (*X, τ*) or (*P, τ*) spaces too.From Eq. 19 one finds that the critical points (*V_c_, S_c_*) are connected by *V_c_* = *εS_c_*/(*μ* + *S_c_*). Since *V_c_* + *S_c_* + *P_c_* = 1 one finds that *P_c_* = 1 − *S_c_* − *εS_c_*/(*μ* + *S_c_*). This means that the reaction velocity *V* is not a monotonically increasing function of *S* as predicted by sQSSA (or decreasing function of *τ* as predicted by the integrated form of sQSSA, Eqs. 5) and other approximations where we have 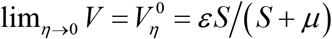. Rather *V* depends on *S* in a turn over manner with a definite maximum at *S_c_*. Particularly we have *V* = 0 for both *S* = 0 and *S* = 1. Expression for *V_c_* suggests that the solutions based on sQSSA will pass through the optimum point (*V_c_, S_c_*) which lie exactly on the trajectory of the integral solution of Eq. 19.

Here one should note that 1 ≥ *S* ≥ *S_c_* in VS space corresponds to the pre-steady state dynamics and *S_c_* ≥ *S* ≥ 0 corresponds to post-steady state dynamics of MMS. Observation of the functional form of the solutions obtained by using sQSSA and associated boundary conditions suggests that it is valid only for the post-steady state dynamics of MMS i.e. *S_c_* ≥ *S* ≥ 0. The expression for *V* in the limit as *ε* → 0 for the pre-steady state regime of MMS 1 ≥ *S* ≥ *S_c_* can be obtained by expanding the right hand side of Eq. 19 in a Macularin series around *ε* = 0 as follows.

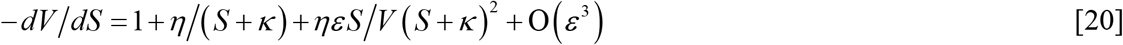

Upon setting *ε* = 0 in this equation and solving the resultant first order ODE for the pre-steady state initial condition as *V* = 0 at *S* = 1 one finds the following expression for *V* which is valid for the pre-steady state range of *S* within 1 ≥ *S* ≥ *S_c_* in the limit *ε* → 0 as follows.

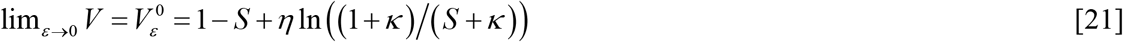

Since *P* = 1 − *S* − *V* one finds from Eq. 21 that 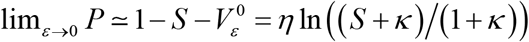 for the pre-steady state range of *S* inside 1 ≥ *S* ≥ *S_c_*. When *ε* → 0 then the approximation given in Eq. 21 will intersect (Fig. 6B) the trajectory of sQSSA at the steady state critical point *S_c_* as follows.

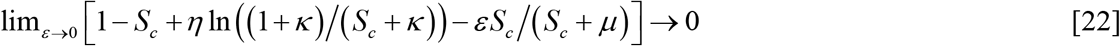

Upon approximating as *εS_c_*/(*S_c_* + *μ*) ≃ *εS_c_*/*μ* one can derive an approximate expression for the steady state critical point *S_c_* by solving Eq. 22 as follows.

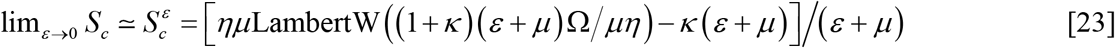

Here we have defined Ω = exp ((*κ*(*ε* + *μ*) + *μ*)/*μη*). Upon substituting the critical value *S_c_* into the expression of *V_c_* in the VS space one can obtain the approximate values of *V_c_* and *P_c_* in the limit *ε* → 0 as follows.

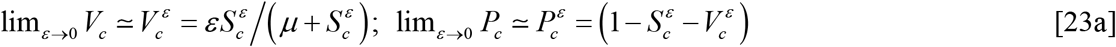

Here Eq. 23a will be valid when *μ* ≫ 1 (which also means that *K_M_* ≫ *s*_0_) since *S_c_* ∈ (0,1). The % error associated with Eqs. 22 and 23 in predicting *S_c_* as 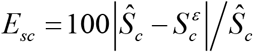 where *Ŝ_c_* is the original value of *S_c_* which can be obtained by numerical integration of Eqs. 3 for a given set of parameters (*κ, η, ε*) for the initial condition (*X, P, S*) = (0, 0, 1) at *τ* = 0 and then identify the time point at which [*dX*/*äτ*] = 0 by numerical differentiation. At this time point we have *S* = *Ŝ_c_*. Here Eqs. 22 and 23 clearly suggest that the stationary reactant assumption i.e. *S_c_* ~ 1 will be valid only when the (1) conditions *ε* → 0 as well as *η* → 0 are strictly true or (2) the conditions *κ* → ∞ is true. Detailed dynamics of MMS in the VS space is demonstrated in Fig. 6B.

### 2.5. Dynamics of MMS in the VP space

From the first equation of Eqs. 12 one can derive the expression for the MMS dynamics in the VP space as follows (Fig. 6A).

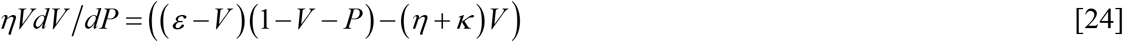

This equation suggests that there also exists an optimum point (*V_c_, P_c_*) in VP space at which[*dV*/*dP*] = 0 i.e. *V* is dependent on *P* in a turnover way with a definite maximum at *P_c_*. Further we have *V* = 0 for both *P* = 0 and *P* = 1 and one needs to select the appropriate root of [*dV*/*dP* ] = 0 which satisfies the post-steady state initial condition as *V* = 0 for *P* = 1 in the VP space. These critical points (*V_c_, P_c_*) are connected by the following relationship which can be obtained by equating the right hand side of Eq. 34 to zero and then solving it for *V*.

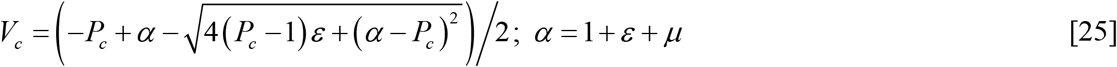

Clearly the post-steady state solution trajectory described by Eq. 16 will pass through the critical points (*V_c_, P_c_*). Here one should note that *P_c_* ≤ *P* ≤ 1 in the VP space corresponds to the post-steady state dynamics and 0 ≤ *P* ≤ *P_c_* corresponds to the pre-steady state dynamics of MMS. The condition required for the validity of Eq. 25 will be obviously [*dV*/*dP*] = 0 (the sQSSA condition *η* → 0 in Eq. 24 will not be applicable here since it eventually drives *V* towards zero) and the trajectory defined by Eq. 25 will pass through (*V_c_, P_c_*). Observation of the functional form of sQSSA solution and the associated initial conditions suggests that it is valid only for the post-steady state dynamics of MMS where *P* ranges in *P_c_* ≤ *P* ≤ 1. Here *V_c_, S_c_* and *P_c_* are precisely the steady state values of the reaction velocity, substrate and product concentrations at which one finds that [*dV*/*dP* = *dV*/*dS* = *dV*/*dτ*] = 0. An approximate expression for the dynamics of MMS in the pre-steady state region of VP space can be derived as follows.

In subsequent sections we denote those solutions obtained from Eq. 7 by setting *ε* → 0 as ***ε*-approximations**. When *ε* → 0 then Eq. 7 reduces to *FdF*/*dU* + (*α*/*η*) *F* − *η*^−3^ ≃ 0 where we have defined *F* = *dU*/*dτ* and *F* is connected to *V* through *V* = *εη*^2^*F*. This follows from the fact that the variable *U* is connected to *P* as *P* = *εη*^2^*U*. The nonlinear ODE corresponding to *F* is a separable one as (*ηF*/(*η*^−2^ − *αF*))*dF* = *dU*. The implicit integral solution of this ODE for the initial condition as *F* = 0 at *U* = 0 will be*U* + *ηF*/*α* + ln(*αFη*^2^ − 1)/*ηα*^2^ = 0. As in case of Eqs. 5 and 13, this equation also can be inverted in terms of Lambert’s W function. Particularly we can obtain the following expression for reaction velocity associated with the pre-steady state dynamics of MMS enzymes in the VP space.

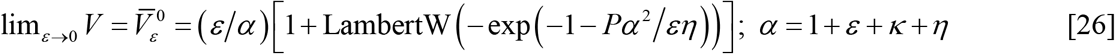

Here we should note that Eq. 26 will be valid only for the pre-steady state region of MMS where *P* ranges inside 0 ≤ *P* ≤ *P_c_*. Clearly the solution trajectory given by Eq. 26 will approximately intersect the solution trajectory (as shown in Fig. 6A) given in Eq. 25 at the steady state critical point (*V_c_, P_c_*) in VP space which will be an exact one in the limit as *ε* → 0. Eq. 26 is one of the central results of this paper which can give accurate description of the pre-steady state dynamics of MMS over wide range of parameter values. Using Eq. 26 one can derive an expression which is more accurate than Eq. 21 for the pre-steady state dynamics of MMS in the VS space as follows. Upon solving the equation 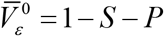 for variable *P* we can obtain the expression for *P* as a function of *S*. Upon substituting this expression for *P* in to the expression for 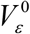 as defined in Eq. 21 one can obtain the following expression for the reaction velocity of MMS enzymes in the pre-steady state region of VS space.

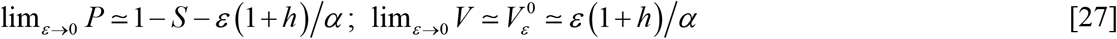

Here *h* is the real root of the following algebraic equation.

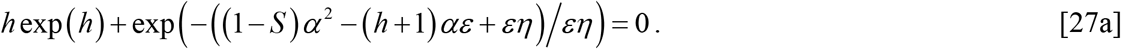

One should note that when *φ* = *ηε* → 0 then Eqs. 27 reduces to an accurate expression for the steady state velocity of MMS similar to 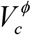 as 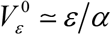 since we have *h* = 0 in such limiting conditions. Detailed dynamics of MMS enzymes in VP space is shown in Fig. 6A.

### 2.6. Average reaction velocity of MMS

Since the reaction velocity *V* is a turnover type functions of *τ, S* and *P*, it will be more appropriate to define an average reaction velocity in VS space especially in the sQSSA limit where *S_c_* ~ 1 for *ε* ~ 0 as follows.

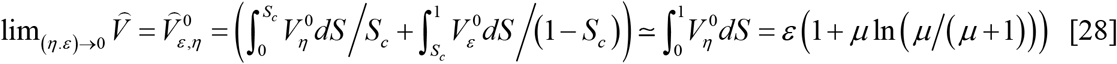

Here 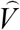 is the real overall average reaction velocity whose expression is unknown, 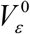 is the approximate reaction velocity in the pre-steady state region of VS space as given in Eq. 21 and 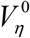 is the approximate reaction velocity associated with the post-steady state region of MMS which resembles sQSSA. In terms of original variables one can express the average reaction velocity associated with sQSSA as follows.

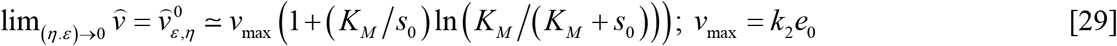

Here we have defined 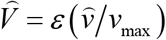 and 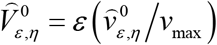. Actually Eq. 29 will be the more appropriate one with respect to *in vitro* experimental procedures where one always measures the reaction velocity averaged over time at different initial substrate concentrations *s*_0_ under sQSSA conditions. The widely used rate equation associated with sQSSA along with the stationary reactant assumption *S_c_* ~ 1 i.e. 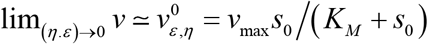 overestimates the reaction rate of MMS enzymes. The error introduced upon using sQSSA to estimate the time averaged reaction velocity can be defined as 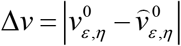. This differential error will be a maximum at *μ* = *μ_m_*. Here *μ_m_* = exp (*y*)/(1 − exp (*y*)) where *y* is the real root of the algebraic equation *y* + exp (2*y*) − 3 exp (*y*) + 2 = 0. Clearly *μ_m_* is not dependent on *ε*. One can obtain this result by solving the equation ∂[Δ*v*]/∂*μ* = 0 for *μ*. Numerical analysis shows that *μ_m_* ≃ 0.46 and the maximum error at this *μ_m_* seems to be Δ*v*_max_ ~ 0.21. It seems that Δ*v*_max_ < 0.01 when *μ* < 10^−3^ or *μ* > 42. Noting that *μ* = *K_M_*/*s*_0_, one can conclude that the differential error Δ*v* can be decreased by setting either *s*_0_ ≫ *K_M_* or *s*_0_ ≪ *K_M_*.

### 2.7. Dynamics of MMS in PS space

Noting the fact that *X* = (1 − *P* − *S*)/*ε*, one can rewrite the rate equations associated with the MMS kinetics in the PS space as follows (Fig. 6C).

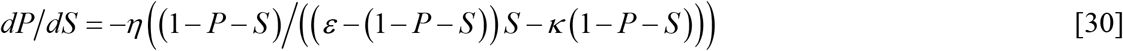

Analytical solution to this nonlinear ODE is not known. Noting the fact that *P* = 1 − *S* − *V*, one can obtain an approximate solution to Eq. 30 especially in the sQSSA limit as 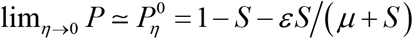. One should note that this formula will work only for the post-steady state dynamics of MMS where *S* ranges from *S_c_* to 0. Here the initial condition for the post-steady state dynamics will be *P* =1 at *S* = 0. The solution that is valid for the pre-steady state region can be obtained as follows. Upon expanding the right hand side of Eq. 30 in a Macularin series around *ε* = 0, one obtains the following series.

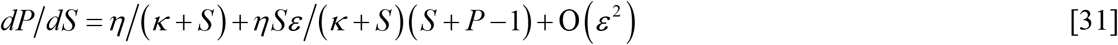

In the limit as *ε* → 0 one finds that *dP*/*dS* ≃ *η*/(*κ* + *S*). Integral solution associated with this first order ODE for the initial condition *P* = 0 at *S* = 1 can be written as follows.

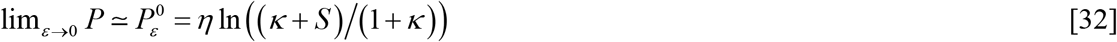

Eq. 32 will be valid only for the pre-steady state dynamics of MMS for the range of substrate *S* from 1 to *S_c_*. One can also derive more accurate expression for the pre-steady state dynamics of MMS in the SP space (variables are interchanged so that here we express *S* as a function of *P*) from Eq. 26 as follows.

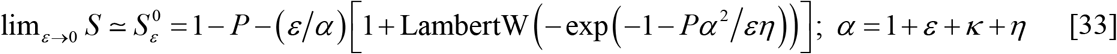

Here one should note that one can also use Eqs. 27 to express *P* as function of *S* in the pre-steady state regime of MMS dynamics in PS space. Clearly the solution sets 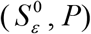 and 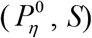 corresponding to the pre- and post-steady state regions of MMS will intersect each other at (*P_c_, S_c_*) in the PS/SP space as shown in Fig. 6C. The approximate of value of *S_c_* can be obtained by numerically solving Eq. 22. Upon substituting the value of *S_c_* in to the expressions for *P* one can obtain the value of *P_c_*.

The expression for 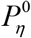 clearly suggests that the approximation *S* ~ 1 − *P* that was used for deriving the integrated rate equation of sQSSA will be valid in the post-steady state region of MMS only when either the limit *ε* → 0 or *μ* → ∞ is true. On the other hand, the expression for 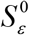 as in Eqs. 33 suggests that the extrapolation of solution obtained from sQSSA through the pre-steady state region will be meaningful only in the limit as *ε* → 0. The overall error (we denote it as **ERIII**) in the approximation used to derive the integrated rate equation can be computed as follows.

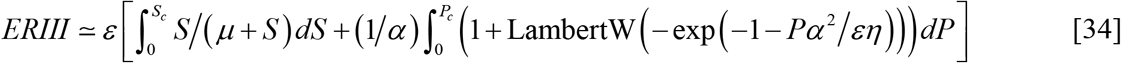

Here **ERIII** is the total area confined between the curves *P* = 1 − *S* and *P* = 1 − *S* − *V* from *S* = 1 to *S* = 0 i.e. 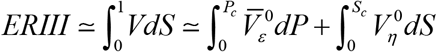. Further Figs. 7A-D show how **ERIII** varies with respect to the parameters (*ε, κ, η*). One can compute *S_c_* and *P_c_* using Eqs. 23 and 23a especially in the limit as *ε* → 0. Figs. 7E-H show how the percentage error *E*_sc_ associated with the prediction of *S_c_* by Eqs. 23 and 23a varies with respect to changes in the parameters (*κ, η, ε*). In deriving this equation we have used the corresponding approximate expression for *V* in the pre- and post-steady state regimes. Eq. 34 suggests that the overall error in the integrated rate equations associated with sQSSA can be minimized by either decreasing *ε* or increasing *μ* which can be achieved by increasing *η* and *κ*. This follows from the facts that *α* = 1 + *ε* + *η* + *κ* and *μ* = *η* + *κ*.

**FIGURE 7:**
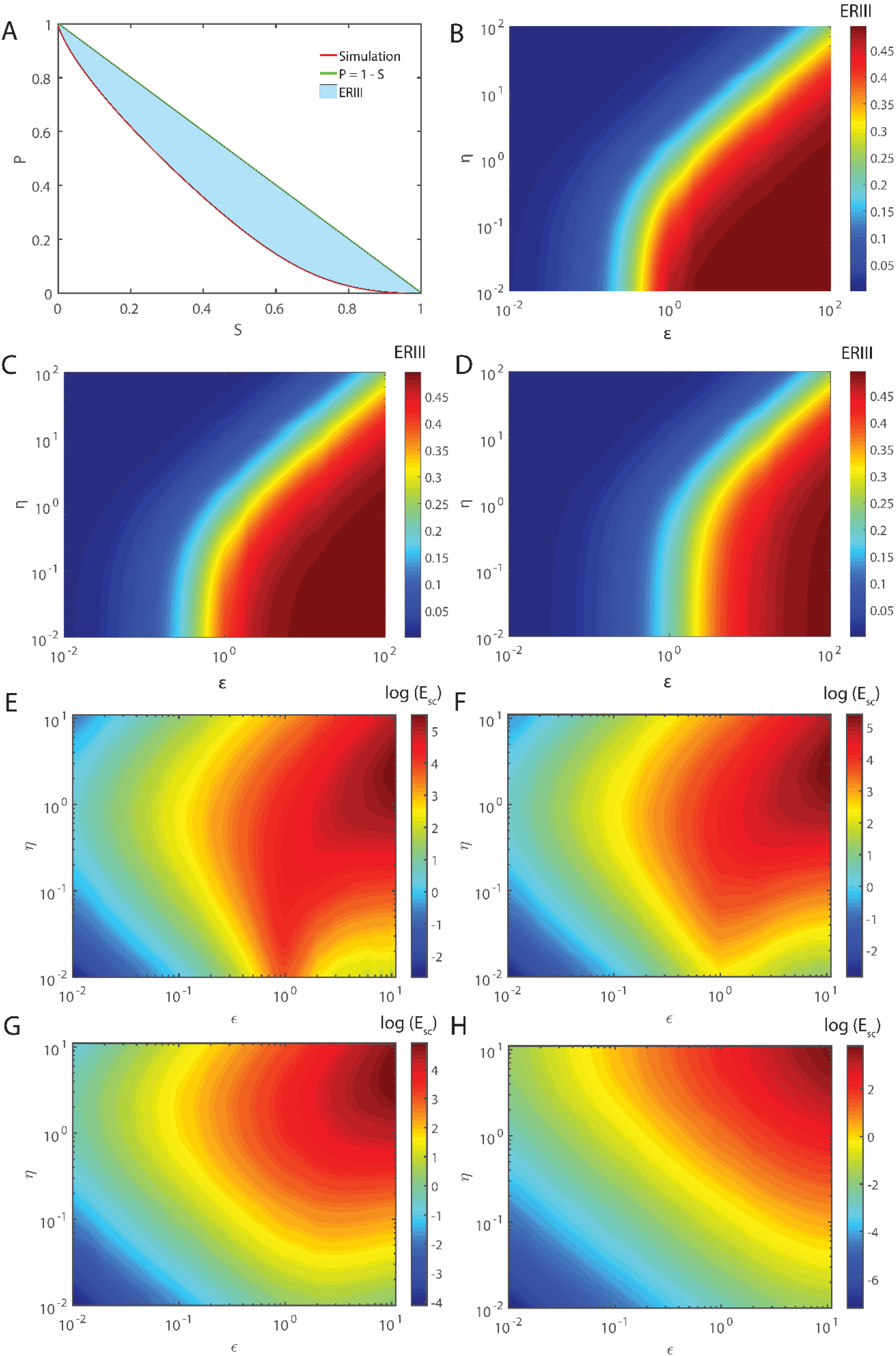
**A-D**. Error in the approximation *P* ~ 1 − *S* (originally we have *P* = 1 − *εX*− *S*) that is used while deriving the integrated rate equations of sQSSA with respect to changes in parameters (*ε, η, κ*). **A**. Here solid red line is the numerical integration of Eq. 30 of PS space with the initial condition *P* = 0 at *S* = 1 and green solid line is the approximation *P* ~ 1 − *S* which requires either the condition *X* ~ 0 or *ε* ~ 0. Error (ERIII) in this approximation is directly proportional to the area confined between these two trajectories. One should note that maximum area = 0.5. Clearly ERIII will be less for low values of *ε* or high values of *η*. High values of *η* ensures less accumulation of *X* and low values of *ε* drives the term *εX* in the definition *P* towards zero. **B**. Here *κ* ~ 0.1. **C**. Here *κ* ~ 1. **D**. Here *κ* ~ 10. **E-G**. Error (*E*_sc_) associated with the approximation i.e. 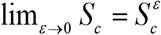 of the substrate concentration at which MMS attains the steady state. Explicitly this percentage error can be written as 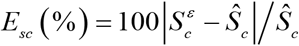 where *Ŝ_c_* is the original value obtained from numerical simulation of Eqs. 3 and 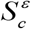 is its approximation obtained by solving Eq. 23. To obtain the original value of *Ŝ_c_* one needs to integrate Eqs. 3 numerically for a given set of parameters (*κ, η, ε*) with the initial conditions as (*X, P, S*) = (0, 0, 1) at *τ* = 0 and then identify the time point at which [*dX*/*dτ*] = 0 by numerical differentiation. At this time point *S* = *Ŝ_c_*. **E**. *κ* = 0.01. **F**. *κ* = 0.1. **G**. *κ* = 1. **H**. Here *κ* = 10.

### 2.8. Dynamics of MMS in the VPS space

Among the dynamical variables (*X, E, S, P*) one should note that the variables (*S, P*) vary with *τ* in a monotonic way and (*X, E*) vary with *τ* in a turn over manner. Our detailed analysis suggests that there exists a critical set of points such that *dX*/*dP* = *dX*/*dS* = *dX*/*dτ* = 0 where (*X, P, S*) = (*X_c_, P_c_, S_c_*) (Fig. 6D). Noting that *V* = *εX*, using Eqs. 19 and 24 one can derive the following partial differential equation corresponding to the dynamics of MMS enzymes in VPS space.

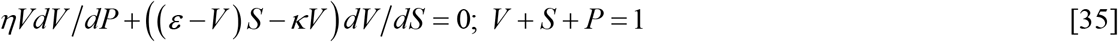

Here the initial condition will be *V* = 0 at *P* = 0 and *S* = 1 for the pre-steady state dynamics. Clearly all the solution trajectories of Eq. 35 will lie on the surface defined by *V* = 1 − *P* − *S*. Though the PDE given in Eq. 35 is not exactly solvable, one can derive an approximate parametric expressions for the dynamic variables (*V, P, S*) from Eq. 8 which is valid only in the limit as *φ* → 0. Noting the expression for 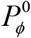 that is given in Eq. 8, one can derive the parametric expressions for 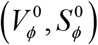 which are all functions of *τ* as follows.

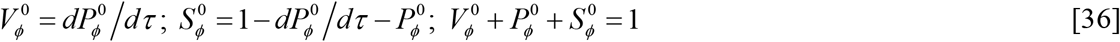

Here one should note that the parametric expressions for 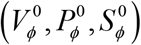 (all are functions of *τ*) satisfy the required initial condition in VPS space as 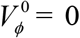 for 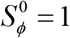 and 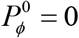. Since there is a *η*^−3^ term in the differential equation corresponding to 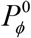 in the limit as *φ* → 0 i.e. left hand side of Eq. 7, Eq. 36 will be close to the solution trajectory of Eq. 35 only in the pre-steady state region of MMS which only requires *ε* → 0. Solution trajectory defined by Eq. 36 deviates from the original solution of Eq. 35 in the post-steady state range of MMS which requires the limiting condition *η* → 0.

One can also derive parametric representations of the solutions to Eq. 36 from Eq. 26 for the pre-steady state region of MMS in VP space as follows.

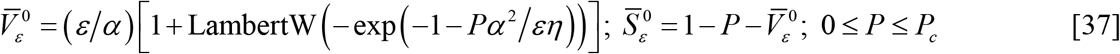

Here 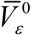 and 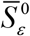 are parametric type functions of *P* and the parametric set of equation in VPS space will be 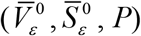. Eq. 37 will be valid for the pre-steady state region of MMS in the VPS space in the limit as *ε* → 0 where the initial conditions for the solution trajectory are 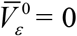 and 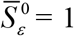 for *P* = 0. Here *P* ranges from *P* = 0 to *P* = *P_c_*. In the same way one can also derive the parametric representations of the solution to Eq. 35 using 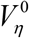 for the post-steady state region of MMS.

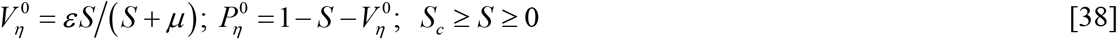

In this equation 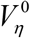 and 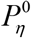 are parametric type functions of *S* and the parametric set of equations in the VPS space will be 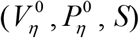. Eq. 38 will be valid for the post-steady state range of MMS in the limit as *η* → 0 where *S* ranges from *S_c_* to 0. Here initial conditions for the trajectories are 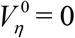 and 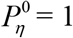 for *S* = 0. One should also note that all those trajectories obtained from both Eqs. 37 and 38 will lie on the plane defined by *V* + *S* + *P* = 1 and they will approximately intersect at the steady state of MMS which occurs at (*V_c_, P_c_, S_c_*) in the VPS space as demonstrated in Fig. 6D. The main results of phase-space dynamics of MMS enzymes in VP, VS, VPS and PS spaces are summarized in Table 2.

**Table 2.**
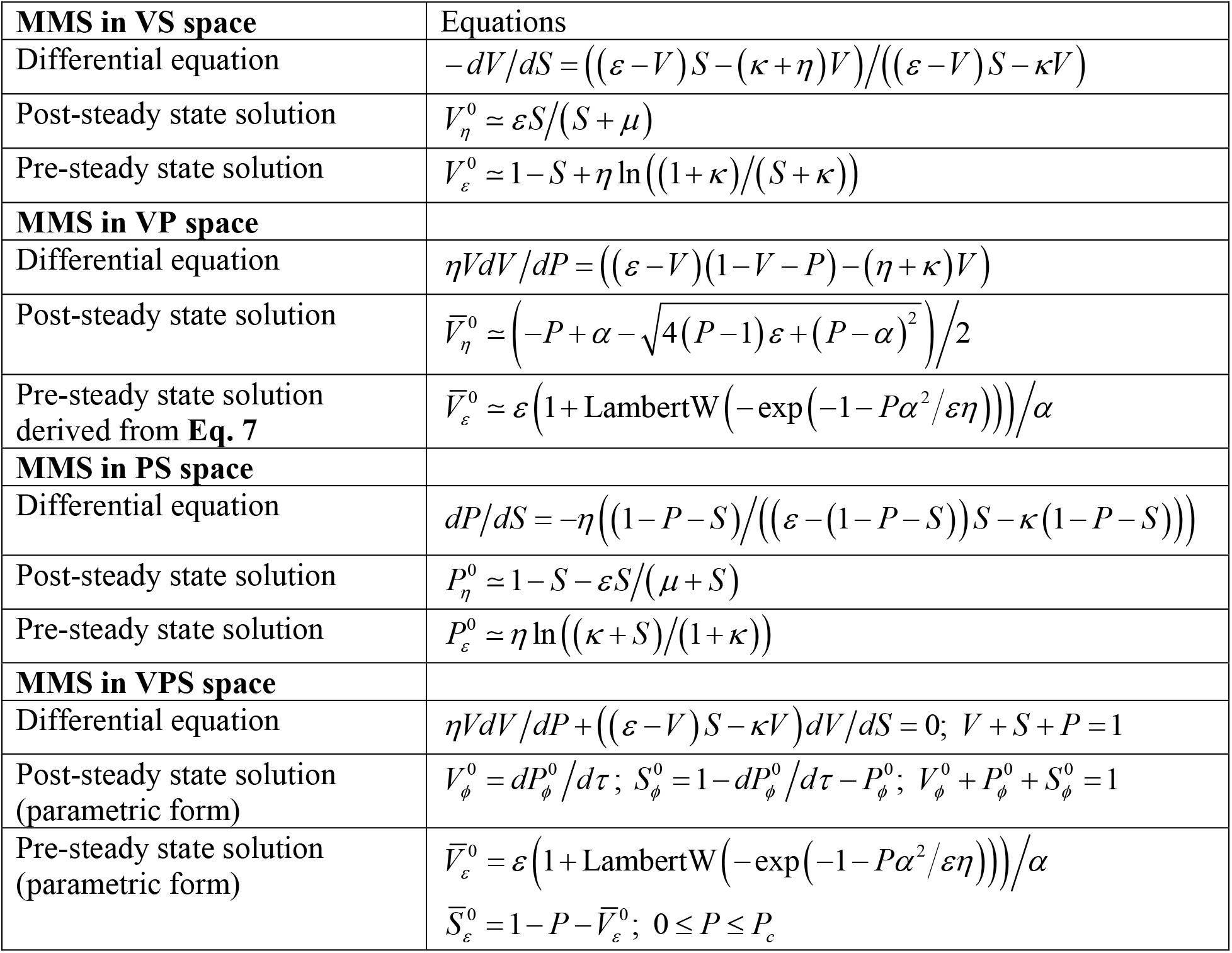
Phase space pre- and post-steady state dynamics of MMS enzymes.

|  |  |
| --- | --- |
| **MMS in VS space** | Equations |
| Differential equation | $-dV/dS = ((\varepsilon - V)S - (\kappa + \eta)V)/((\varepsilon - V)S - \kappa V)$ |
| Post-steady state solution | $V_\eta^0 \simeq \varepsilon S/(S + \mu)$ |
| Pre-steady state solution | $V_\varepsilon^0 \simeq 1 - S + \eta \ln((1 + \kappa)/(S + \kappa))$ |
| **MMS in VP space** |  |
| Differential equation | $\eta V dV/dP = ((\varepsilon - V)(1 - V - P) - (\eta + \kappa)V)$ |
| Post-steady state solution | $\bar{V}_\eta^0 \simeq \left( -P + \alpha - \sqrt{4(P - 1)\varepsilon + (P - \alpha)^2} \right) / 2$ |
| Pre-steady state solution derived from **Eq. 7** | $\bar{V}_\varepsilon^0 \simeq \varepsilon \left( 1 + \text{LambertW} \left( -\exp(-1 - P\alpha^2/\varepsilon\eta) \right) \right) / \alpha$ |
| **MMS in PS space** |  |
| Differential equation | $dP/dS = -\eta \left( (1 - P - S) / ((\varepsilon - (1 - P - S))S - \kappa(1 - P - S)) \right)$ |
| Post-steady state solution | $P_\eta^0 \simeq 1 - S - \varepsilon S/(\mu + S)$ |
| Pre-steady state solution | $P_\varepsilon^0 \simeq \eta \ln((\kappa + S)/(1 + \kappa))$ |
| **MMS in VPS space** |  |
| Differential equation | $\eta V dV/dP + ((\varepsilon - V)S - \kappa V) dV/dS = 0; V + S + P = 1$ |
| Post-steady state solution (parametric form) | $V_\phi^0 = dP_\phi^0/d\tau; S_\phi^0 = 1 - dP_\phi^0/d\tau - P_\phi^0; V_\phi^0 + P_\phi^0 + S_\phi^0 = 1$ |
| Pre-steady state solution (parametric form) | $\bar{V}_\varepsilon^0 = \varepsilon \left( 1 + \text{LambertW} \left( -\exp(-1 - P\alpha^2/\varepsilon\eta) \right) \right) / \alpha$<br>$\bar{S}_\varepsilon^0 = 1 - P - \bar{V}_\varepsilon^0; 0 \leq P \leq P_c$ |

### 2.9. Temporal pre- and post-steady state dynamics of MMS

One can derive the expressions for the time dependent reaction velocity, substrate depletion and product evolution as follows. When *ε* → 0 then Eq. 7 reduces to a linear ODE as *dH*/*dτ* + (*α*/*η*)*H* − *η*^−3^ ≃ 0 where we have defined *H* = *dU*/*dτ* and *H* is connected to the reaction velocity *V* as *V* = *dP*/*dτ* = *εη*^2^*H*. The integral solution to this ODE for the initial condition as *H* = 0 at *τ* = 0 can be written in terms of *P* and *V* as follows.

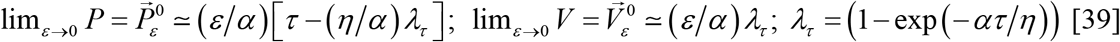

Noting that *S* = 1 − *P* − *V*, one can derive the following expression for the temporal substrate depletion curve which is valid in the pre-steady state region of MMS.

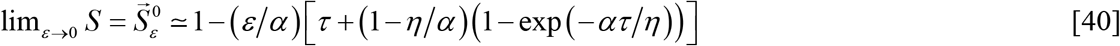

Here forward arrow over *S* denotes the pre-steady state dynamics. Eq. 39 clearly suggests the pre-steady state timescale as lim_*ε*→0_ *t_c_* = (*η*/*k*_2_*α*) in line with Eqs. 9. In terms of the original variables one finds that lim_*ε*→0_ *t_c_* = (1/*k*_1_ (*s*_0_ + *e*_0_ + *K_M_*)). For the purpose of convenience we call Eqs. 39 and 40 as ***ε*-approximations**. The approximation given by Eqs. 5 (which actually requires (*η, ε*) → 0) accurately fits the substrate depletion data well only in the post-steady state region and Eq. 40 fits only the pre-steady state region of MMS i.e. particularly one should note that Eq. 40 will be valid only for the range of *τ* in (0, *τ_c_*). These results are summarized in Figs. 8A-B. In the limit as *ε* → 0, Eqs. 39 and 40 seems to predict the burst phase of MMS enzymes very nicely as shown in Figs. 8A and 8C.

**FIGURE 8:**
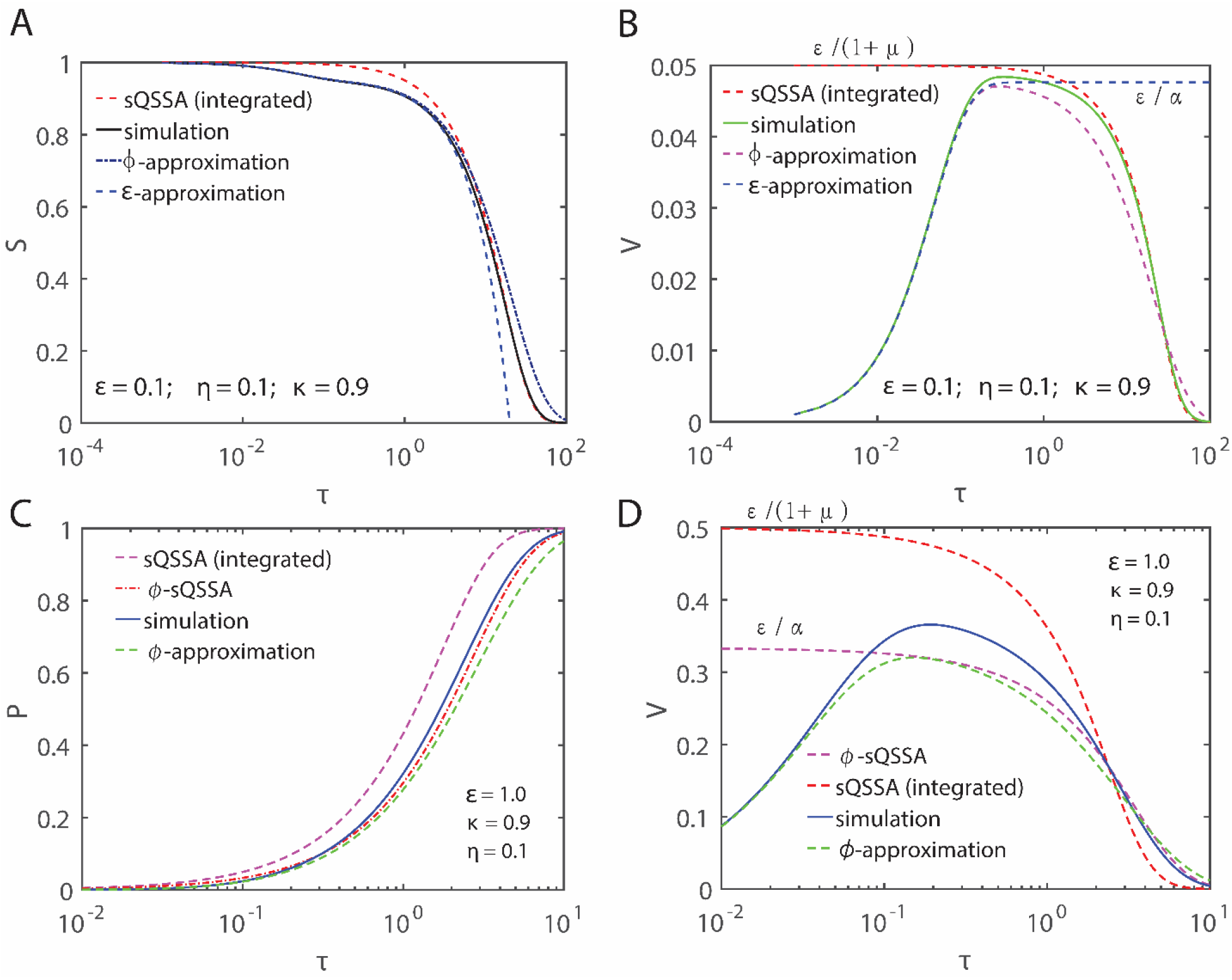
Approximations to the pre- and post-steady state dynamics of MMS. **A**. Here red dotted line is the approximation by integrated form of sQSSA (Eqs. 5). Black solid line is the numerical simulation using Eqs. 3. We computed as 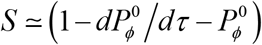 using Eq. 8 for the *φ*-approximation of *S* and used Eqs. 40 for the *ε*-approximation. Clearly *ε*-approximation works only in the pre-steady state region. On the other hand, integrated form of sQSSA will be accurate only in the post-steady state region. Though φ-approximation works over the entire range well, there is a slight deviation from the simulation especially in the post-steady state region. **B**. Simulation results show that *V* = 0 for both *τ* → 0 and *τ* → ∞. Although the integrated form of sQSSA (where it is assumed that *V* ≃ −*dS*/*dτ*) works very well in the (*S, τ*) space (with slight deviation from the simulation in the pre-steady state region), it fails to predict the pre-steady state region of MMS dynamics in (*V, τ*) space. On the other hand *ε*-approximation fails to model the post-steady state region of MMS dynamics in (*V, τ*) space. Although *φ*-approximation works very well over the entire range of time, there is a slight deviation from the simulation result especially in the post-steady state region of MMS dynamics in (*V, τ*) space. **C**. Comparison of integrated from of sQSSA (Eqs. 5, we used the approximation *P* ~ 1 − *S* to calculate *P*) and *φ*-sQSSA for the product evolution curve as in Eq. 41 at *ε* = 1.0. **D**. Velocity space comparison of approximations presented in **C. C** and **D** clearly suggest that *φ*-sQSSA is more accurate than integrated form of sQSSA over the entire range of *ε*.

One can use Eqs. 13 to model the post-steady state product evolution curve. An expression which is more accurate than Eqs. 13 can be derived by retaining *ε* terms in Eqs. 12 as follows. Upon setting *ε* → 0 in the first equation of Eqs. 12 and noting that *X* = [*dP*/*dτ*]/*ε* one can obtain ((*α* − *P*)/*ε*)[*dP*/*dτ*] + *P* − 1 ≃ 0. This is a separable nonlinear ODE similar to the equation of sQSSA as ((*α* − *P*)/*ε*(1 − *P*)) *dP* = *dτ*. Unlike in case of Eqs. 13, here we have ignored only those higher order polynomial or derivative terms which multiply *ε*. Upon solving this nonlinear ODE for the initial condition as *P* = 0 at *τ* = 0, one can express the solution in terms of Lambert’s W function as follows.

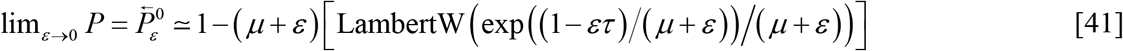

Upon comparing this equation with Eqs. 5 and 13 one can conclude that only in the limit as *ε* → 0 we have 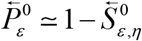. However those *μ* terms in Eq. 13 needs to be replaced with *μ* + *ε* for an accurate fitting results (Tzafriri, 2003) which is evident from Figs. 8C-D. Tzafriri in Ref. (Tzafriri, 2003) has derived an expression similar to Eq. 41 (Eq. 55 in Ref. (Tzafriri, 2003)) based on tQSSA arguments. One should note that the approximation given by 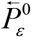 will be valid even though when the strict sQSSA condition *η* → 0 is not true. While deriving Eq. 41 we have not used any assumptions like *dP*/*dτ* ≃ −*dS*/*dτ* which were actually the basis for deriving Eqs. 5 as well as Tzafriri model. Here one should note that Eqs. 41 is valid only for the post-steady state dynamics of MMS and it also has the flaws of Eqs. 5 and 13 in the velocity-time space (*V, τ*) which is clearly evident from the limiting condition 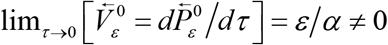.

From Eq. 41 one can derive approximate expressions for the temporal evolution of substrate and velocity (*S, V*) corresponding to the post-steady state regime of MMS dynamics in the limit *ε* → 0 as 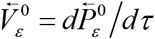 and 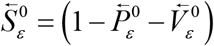. For the purpose of convenience we can call the approximation given in Eq. 41 as ***φ*-sQSSA**. Detailed numerical simulations in Figs. 8C-D suggest that *φ*-sQSSA is more accurate than the integrated forms of sQSSA given by Eqs. 5 and 13. Particularly Eq. 41 predicts the (*P, τ*) progress curve very well even under the conditions like *ε* ≥ 1.

### 2.10. Error associated with the integrated sQSSA methods

Since Eqs. 39 and 40 fits the pre-steady state data well in the limit as *ε* → 0, one can compute the approximate error (Ψ) introduced upon using the integrated version of sQSSA i.e. Eqs. 5 over pre-steady state region as follows.

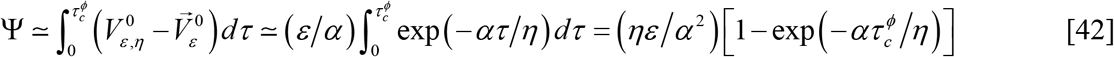

The pre-steady state reaction velocity associated with the integrated form of sQSSA can be calculated from Eqs. 5 as 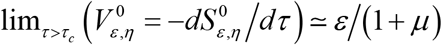 which is approximately equal to *ε*/*α*. Since we have 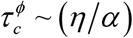 which is approximately the time required to attain the steady state, one finds that Ψ ≃ (*ηε*/*α*^2^)[1 − exp(−1)] ≃ 0.63 (*ηε*/*α*^2^). This error level will be a maximum at some critical value *η_m_* where *η_m_* ≃ (1 + *ε* + *κ*). This can be obtained by substituting *α* = 1 + *ε* + *κ* + *η* in to the expression for Ψ and then solving the algebraic equation [∂Ψ/∂*η*] = 0 for *η*. Upon substituting this value of *η_m_* back in to Eq. 42 one can obtain an approximate estimate of the maximum possible error associated with the integrated form of sQSSA Eqs. 5 as Ψ_max_ ≃ 0.16(*ε*/*η_m_*). This further suggests that Ψ_max_ asymptotically increases with *ε* and decreases with *κ*.

Instead one can also define a critical value *ε_m_* where *ε_m_* ≃ (1 + *η* + *κ*) at which the overall error Ψ is a maximum. This can be obtained by solving the equation [∂Ψ/∂*ε*] = 0 for *ε*. The maximum error in this particular case will be Ψ_max_ ≃ 0.16 (*η*/*ε_m_*). These results also suggest the stringent condition that is required for attaining a minimum amount of error in the sQSSA methods as *ε* ≪ *ε_m_*. One can write this condition explicitly as *ε* ≪ (1 + *μ*) where *μ* = *η* + *κ*. This also means that (*ε*/*μ*)≪((1 + *μ*)/*μ*). In terms of the original variables one can write this as (*e*_0_/*K_M_*)Ȫ(1 + *s*_0_/*K_M_*) which was derived earlier as a stringent condition for the validity of sQSSA using different set of arguments (Hanson and Schnell, 2008; Segel, 1988; Stroberg and Schnell, 2016). Our results further suggest that the overall error Ψ associated with the pre-steady state region of integrated form of sQSSA can also be minimized by setting the condition that *η* ≪ *η_m_* i.e. (*α*/*η*) ≫ 2. We can summarize the generalized conditions required for minimizing the error in the integrated sQSSA as follows.

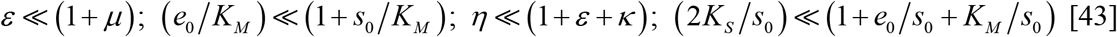

Here the second condition in this equation also means that 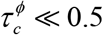 or 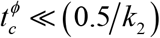. In terms of the original variables of MMS one can rewrite this inequality as (2*K_S_*) ≪ (*s*_0_ + *e*_0_ + *K_M_*). Here one should note that the overall error in sQSSA cannot be decreased by setting *ε* ≫ *ε_m_* or *η* ≫ *η_m_* since these conditions will eventually invalidate sQSSA.

## 3. Results and Discussion

### 3.1. Conditions for the validity of sQSSA

Earlier studies (Borghans et al., 1996) suggested that the sQSSA models used to fit experimental data (where the reactant stationary assumption *S* ~ 1 is used) will be valid only when (**a**) there is a significant timescale separation between pre- and post-steady state dynamics and (**b**) substrate concentration is much higher than enzyme concentration. We first define these conditions in line with the parameters (*η, κ, ε*) of MMS dynamics.

*Condition* **C1**. The timescale separation (Borghans et al., 1996) required for the validity of sQSSA can be well described by the parameter *η* = *k*_2_/*k*_1_*s*_0_. Here 1/*k_2_s_0_* is the timescale associated with the pre-steady state dynamics of MMS and 1/*k_2_* is the timescale associated with the post-steady state dynamics. Clearly *η* should be close to zero for the validity of sQSSA.

*Condition* **C2**. The ratio of initial concentrations of enzyme to substrate as *ε* = *e*_0_/*s*_0_ should be close to zero. When **C1** is true, then the accumulated steady-state concentrations of *x* and *p* at the end of the pre-steady state will be much lesser than *s*_0_ and one can approximate as *s* ≃ *s*_0_ as in case of reactant stationary approximation (Stroberg and Schnell, 2016). In the dimensionless form one can write the reactant stationary approximation as *S* = (1 − *εX* − *P*) ~ 1.

As we have shown in section 2.2, the timescale separation ratio is strongly dependent on (*ε, η, κ*) i.e. 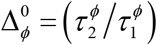. Particularly we have 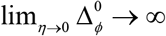 and 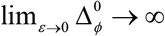. Consequently conditions **C1** and **C2** are mandatory for enhancing the timescale separation ratio. Bajzer and Strehler (Bajzer and Strehler, 2012) addressed the scenario where **C1** is true but **C2** is not met i.e. *ε* ≥ 1. Borghans et.al in Ref. (Borghans et al., 1996) and Tazfriri in Ref (Tzafriri, 2003) addressed similar situations in their earlier treatments on tQSSA. However they have not performed detailed comparative error analysis as in the works of Bajzer and Strehler in Ref. (Bajzer and Strehler, 2012). In this situation, when **C1** is true then the system attains the steady state quickly with most of the substrate molecules bound with the excess enzyme. Further **C1** ensures slow formation of the product which leads to the approximation *s* ≃ (*s*_0_ − *x*) used to derive 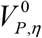 (in section 2.3). When **C1** is not true then there will be a rapid accumulation of the product molecules and this approximation will not be valid. The error analysis results given by Bajzer and Strehler in Ref. (Bajzer and Strehler, 2012) validate these arguments. In this paper we propose an alternative but effective reformulation of sQSSA. We suggest the following generalized condition **C3**.

*Condition* **C3**. *φ* = *εη* = 0 (*φ*-approximation) This limit can be easily achieved even under slight violation of any one of the conditions **C1** and **C2**. This is to say, when **C1** is strongly true then **C2** can be slightly relaxed and the *vice versa*.

### 3.2. Error level analysis of QSSAs and steady state φ-approximations

For detailed comparison of the error levels of various steady state approximations we considered the expressions associated with sQSSA (conditions **C1** and **C2** are strictly true), Bajzer and Strehler model (**C1** is true and **C2** is relaxed) and *φ*-approximation (**C1** is true and **C2** is relaxed and the *vice versa*). The true value of the steady state reaction velocity was calculated by numerically integrating the reduced version of Eqs. 3 using the following Euler iterative scheme (Abramowitz and Stegun, 1965).

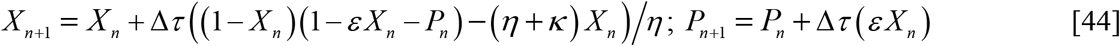

Here the initial conditions are (*X_0_, P_0_*) = (0, 0) and Δ*τ* ~ 10^−5^. Error level analysis were done over a wide range of parameters (*ε, η, κ*). The original steady state reaction velocity was obtained by numerically differentiating the temporal trajectory of *X* with respect to *τ* and subsequently identifying the location of change in the sign of [*dX*/*dτ*] where *X_c_* occurs. At this point the steady state velocity will be *V_c_* = *εX_c_* where [*dV*/*dP* = *dV*/*dS* = *dV*/*dτ*] = 0 and obviously we have *S_c_* + *V_c_* + *P_c_* = 1. We follow the error level metrics proposed by Bajzer and Strehler in Ref. (Bajzer and Strehler, 2012). We defined the error (%) as 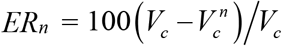 where *V_c_* is the true steady state velocity obtained by numerical integration as in Eq. 44 and 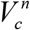 is the steady state velocity obtained from various approximations where *n* = **1** corresponds to sQSSA, *n* = **2** represents Bajzer and Strehler model (Bajzer and Strehler, 2012) and **3-4** represent various forms of steady state *φ*-approximations. The expressions for the steady state reaction velocity under different approximations can be written along with the conditions of validity in terms of (*η,ε,κ*) as follows. Here 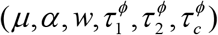 are defined as in Eqs. 8-10.

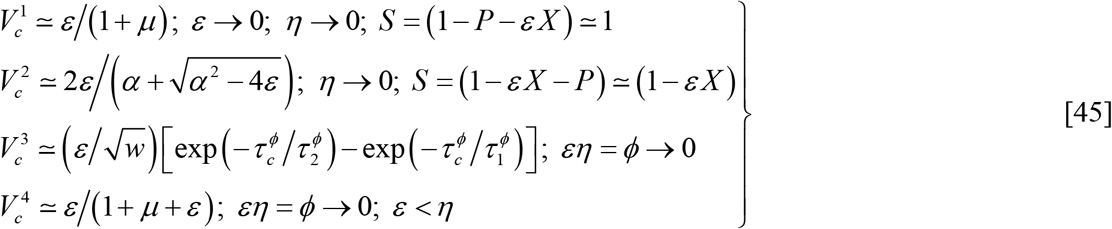

The detailed error level analysis results are summarized in Figs. 9. One can conclude from Figs. 9A1-4 that sQSSA with stationary reactant assumption strictly requires both conditions **C1** and **C2**. When **C2** is met (*ε* ≪ 1) then the error in sQSSA upon violating **C1** shows a maximum with respect to changes in *η* at constant *ε*. The value of *η* at which maximum error occurs seems to increase as *κ* increases. When *ε* ≃ 0.1 and *κ* < 0.1 then the error associated with sQSSA seems to vary from 1 to 10% with respect to the variation of *η* in the range (10^−2^, 10^2^). Clearly sQSSA is still a good approximation for practical purposes when **C2** is strictly met at least with *ε* ≤ 0.1. However, when condition **C2** is not met then irrespective of the validity of **C1** the error levels shoots beyond >1000%.

**FIGURE 9.**
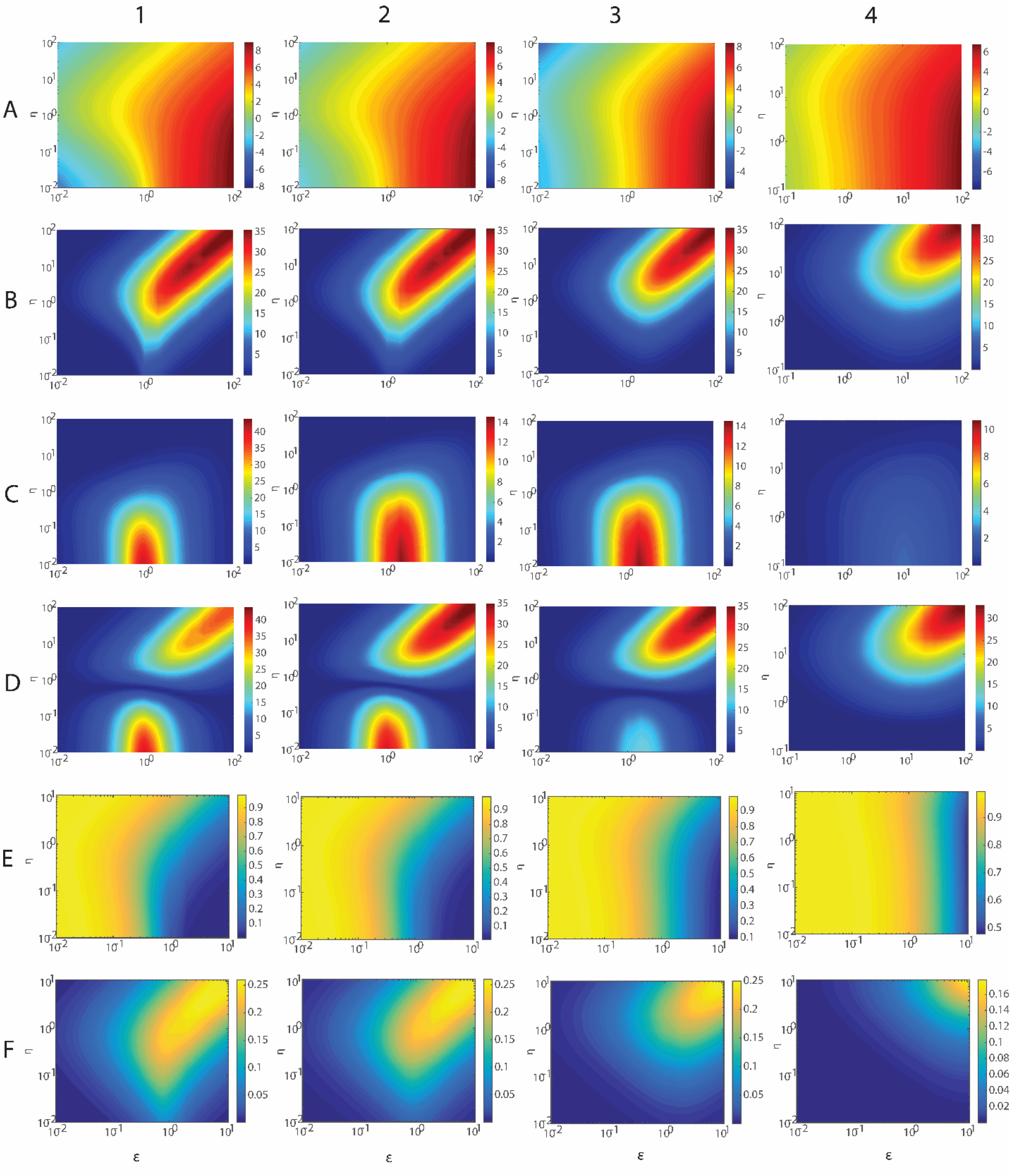
Error level analysis of various approximations of the steady state reaction velocity of single-substrate enzyme catalyzed reactions following MMS. The parameters *η* and *ε* was iterated from 10^−2^ to 10^2^. The parameter *κ* was iterated across 10^−2^, 10^−1^, 1 and 10 (**Columns 1, 2, 3** and **4** respectively). For a given set of (*η, ε, κ*), Eqs. 3 was numerically integrated from τ = 0 to *τ* = 250. Then the steady state reaction velocity (*V_c_*) was obtained by numerically differentiating the temporal trajectory of *X*(since *V* = *εX*) with respect to *τ* and subsequently identifying the location of change in the sign of [*dV*/*dτ*]. We defined the error level 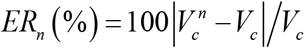 where *V_c_* = *εX_c_* is the steady state reaction velocity obtained by numerical integration of Eqs. 3 and 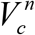 is the steady state reaction velocity obtained from various approximations where *n* = 1, 2, 3, and 4 corresponding to various approximations given in Eqs. 45. Here (**A**) loge (***ER_1_***) is the error in sQSSA with reactant stationary assumption (**B**) ***ER_2_***. (**C**) ***ER_3_*** is the error associated with the φ-approximation. (**D**) ***ER_4_*** is the simplified versions of *ε*- or *φ*-approximation. *ER* values are measured in %. One should note that the colour map scale is different for different panel. **E-F**. Variation of steady state values of substrate (**E**) and product (**F**) (*S_c_, P_c_*) with respect to changes in parameters (*ε, η, κ*). To obtain (*S_c_, P_c_*) one needs to integrate Eqs. 3 numerically for a given set of parameters (*κ, η, ε*) with the initial conditions as (*X, P, S*) = (0, 0, 1) at *τ* = 0 and then identify the time point at which [*dX*/*dτ*] = 0 by numerical differentiation. At this time point (*S, P*) = (*S_c_, P_c_*).

The error levels associated with the Bajzer and Strehler model (Bajzer and Strehler, 2012) is shown in Figs. 9B1-4. Clearly this approximation works much better than sQSSA. The overall error varies from 0-35% for the entire parameter space explored in this paper. Lowest error values are shown up when (1) conditions **C1** and **C2** are strictly met and (2) violation of condition **C1** when **C2** is true and the vice versa. Here maximum error values are shown up when both conditions **C1** and **C2** are violated.

The error associated with the full form of *φ*-approximation 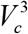 is shown in Figs. 9C1-4. Clearly the *φ*-approximation 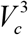 works much better than 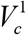 and 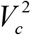 of Eqs. 45 over the entire parameter space explored in this paper except for the region confined by *ε* in (10^−1^, 10^1^) for *η* < 1 and *κ* < 0.1 where the error shoots up to 35-40%. It seems that 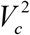 and 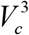 are complementary to each other in terms of error levels over parameter space i.e. 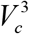 works much better in those parameter regions where 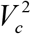 fails and the vice versa. Apparently the error levels are maximum around *ε* = 1 i.e. equal amount of enzyme and substrate in the system. For the range of *κ* in (10^−1^, 1), the overall error level for the entire parameter space varies from 0-15%. When *κ* = 10 then the error level for the entire parameter space was < 2% which is a remarkable observation.

The error diagrams associated with 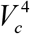 of *φ*-approximation (Eqs. 45) are shown in Figs. 9D1-4. Clearly this approximation is much better than 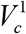 of sQSSA. Here the error levels for the entire parameter space varies from 1-40%. However it has the disadvantages of both 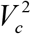 and 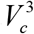. On the other hand, from the practical applications point of view 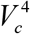 is much better than 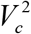 especially when both the conditions **C1** and **C2** are slightly violated.

Our detailed analysis suggested that the error levels of various approximations vary with the parameters (*η, ε, κ*) in a dissimilar manner. Figs. 9E1-4 clearly suggest that the error levels associated with 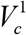 of sQSSA is inversely correlated with the steady state value of substrate *S_c_* which is approximated as *S_c_* ~ 1 in the reactant stationary assumption used to derive the expression for 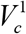. Figs. 9F1-4 clearly suggest that the error levels associated with 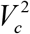 is positively correlated with the steady state value of product *P_c_* which is approximated as *P_c_* ~ 0 in the Bajzer and Strehler model. In most of the experimental situations on single-substrate enzyme reactions, only *ε* will be the known quantity. Therefore based on the initial value of the enzyme to substrate ratio, one needs to decide on the type of approximation formula to be used for the steady state data analysis. To start with one can use 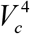 of *φ*-approximation rather than 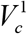 of sQSSA. Here we should note that the expression corresponding to 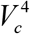 will give a slightly different form under Lineweaver-Burk representation (Lineweaver and Burk, 1934).

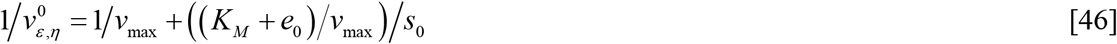

This equation suggests that 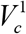 of sQSSA (Eqs. 45) always overestimates the Michaelis-Menten constant *K_M_* when **C2** is violated i.e. using sQSSA methods one always obtains *K_M_* + *e_0_* instead of obtaining the original values of *K_M_*. This is also evident from Eq. 41. Although 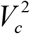 and 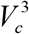 are quiet accurate in this context, they require sophisticated nonlinear least square fitting procedures to obtain the steady state enzyme parameters. Since different approximations work efficiently in different parametric regions, it is mandatory to analyze the steady state experimental data with a combination of approximation formulae. For example one can use a combination of approximations given by 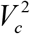 and 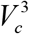 of Eqs. 45 to fit the same dataset. That is to say, we use two different types of approximations to analyze the same dataset. Consistent outcomes of parametric estimates from both these analyses will in turn ensure the correctness of the obtained enzyme parameters. Inconsistent outcomes will give us an idea about the probable range of parameter space associated with the error values.

### 3.3. Error associated with the progress curve analysis

Various kinetic and steady state parameters associated with the MMS enzyme can also be extracted from the progress curve analysis (Duggleby, 1986; Duggleby and Morrison, 1978; Ellis and Duggleby, 1978; Zavrel et al., 2010). Here we can consider both the normalized substrate depletion as well as normalized product evolution data i.e. dataset on (*S, t*) or (*P, t*) where time *t* has the dimension of seconds. For a comparative study, we generated such datasets with various noise levels using the following Euler scheme to integrate Eqs. 3 (slightly modified version of Eqs. 44).

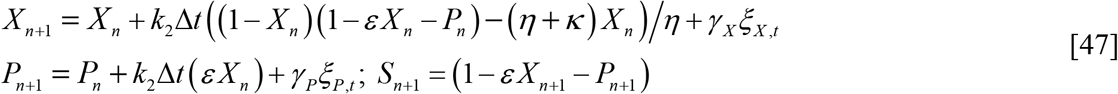

Here *ξ_X, t_* and *ξ_P, t_* are delta-correlated Gaussian white noises with zero mean and unit variance with the following cross covariance properties.

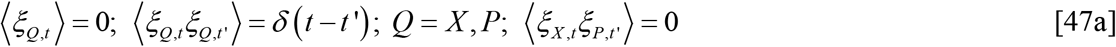

Here *γ_X_* and *γ_P_* are the parameters which control the noise levels of *X* and *P* in the generated datasets. For the purpose of comparison we considered the following data fitting methods.

#### 3.3.1. Case I: Full model or direct nonlinear least square fitting

One can directly fit the normalized substrate depletion or product evolution data to the following differential equation using the standard Marquardt-Levenberg algorithm (Marquardt, 1963).

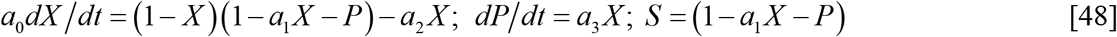

This equation clearly suggests that the value of the initial substrate concentrations (*s*_0_) is necessary to obtain the MMS parameters from the normalized substrate depletion or product evolution datasets i.e. (*P, X, S, t*). Various nonlinear least square fitting parameters associated with Eqs. 48 are defined along with their standard errors (*δ_s_*) at a given confidence level (*θ*, we set *θ* ≃ 0.95 for all the nonlinear least square fittings of this paper) as follows.

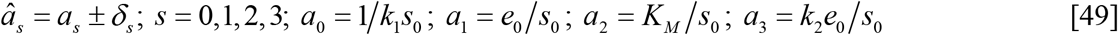

Upon obtaining the best fit values *â_s_* and then following the error propagation theory (Sacks et al., 1989) one can compute the best fit values of *K_M_* and *v*_max_ along with their standard errors as follows.

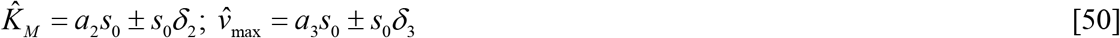

Clearly this fitting procedure requires only the initial substrate concentration as additional input. The main disadvantage of Case I is that a numerical integration of Eqs. 48 is required for each iteration of Marquardt-Levenberg algorithm. It is not possible to obtain the estimate about various timescales associated with the dynamics of MMS using this method.

#### 3.3.2. Case II: Integrated version of sQSSA

When the conditions (**C1** and **C2**) required for sQSSA are strictly met then one can consider the following integrated version of sQSSA.

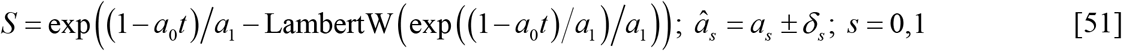

Various nonlinear least square fitting parameters associated with this normalized substrate depletion equation are defined along with their standard errors as follows.

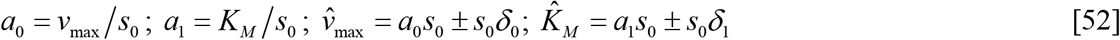

As in Case I, this fitting procedure too requires only the initial substrate concentration *s*_0_ as an additional input along with the normalized substrate depletion data and it is not possible to estimate various timescales associated with the dynamics of MMS using this method. Further only the substrate depletion data can be used for the fitting purposes since there is no way of deriving expression for the product evolution data from Eq. 51. To fit the normalized product evolution dataset (*P, t*) we can use either Eq. 13 or the *φ*-sQSSA given by Eq. 41 as follows.

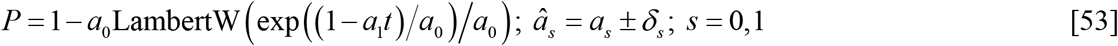

Various nonlinear least square fit parameters associated with this normalized product evolution equation are defined along with their standard errors as follows.

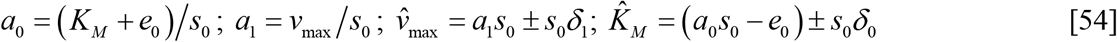

The main advantages of Case II are (1) it has only two fit parameters and (2) one can compute both *K_M_* and *v*_max_ directly from the nonlinear least square fitting results as in Case I. However Eqs. 41 and 54 clearly suggest that the value of *K_M_* obtained from Eqs. 5 (when it is used over product evolution data using *P* = 1 - *S* relationship) is actually an overestimate as *K_M_* + *e_0_*. To get an accurate estimate of *K_M_*, one needs to subtract *e_0_* from 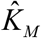 of Eqs. 5 as in Eqs. 54 when it is used over product evolution data.

#### 3.3.3. Case III: *φ*-approximation

When the conditions (**C3**) associated with the *φ*-approximation are true, then both the substrate depletion and product evolution data (or reaction velocity data) i.e. ([*S, P, V*], *t*) can be used to fit with a two exponential function with four distinct parameters as follows.

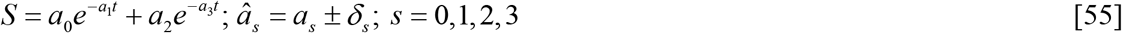

This equation can be derived from the expression for 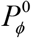 as given in Eq. 8 using the relationship 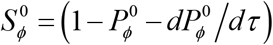. Various parameters associated with this nonlinear least square fit function are defined as in Eq. 8 as follows.

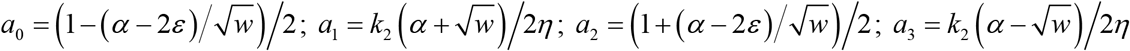

Using these nonlinear least square fit parameters one can obtain the *φ*-approximations of the various timescales associated with the pre- and post-steady state dynamics of MMS along with their standard errors (at a given confidence level, *θ*) as follows.

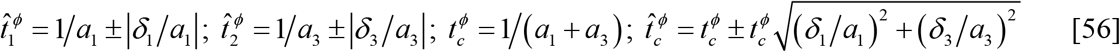

However the main disadvantage of this method is that there is no direct way of obtaining *K_M_* and *v*_max_ from this analysis as in Cases I and II. Using a combination of analyses based on Cases I and III or Cases II and III one can completely characterize the temporal as well as steady state dynamics of a single substrate MMS enzyme. Here one should note that Eqs. 52 will have the disadvantages of multi exponential fitting methods such as degeneracy in the pre-exponential coefficients etc. Accuracy of two exponential fit can be enhanced by using the reaction velocity data (*V, t*) rather than the substrate depletion (*S, t*) or product evolution data (*P, t*). Here one should note that the product evolution data can transformed into velocity evolution data by numerical differentiation (*V, t*) = (*dP*/*dt, t*). The double exponential fit function associated with the *φ*-approximation of the reaction velocity data in terms of original variables can be written as follows.

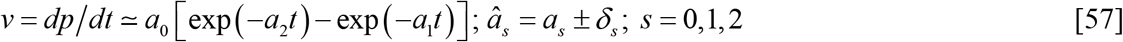

Here various fit parameters along with their standard errors (computed with *θ* confidence level) are defined as in Eqs. 53.

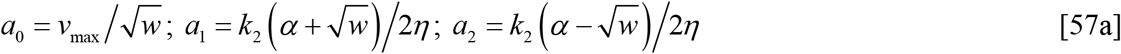

Using the best fit values of the parameters (*â_s_*) one can finally obtain various timescale components associated with the dynamics of MMS as follows.

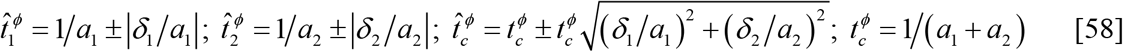

This equation is more efficient in accurately extracting various timescale components from the time derivative of the product evolution data. Because clear demarcation of the pre- and post-steady state is possible over the (*v, t*) data rather than on (*S, t*) or (*P, t*) datasets.

#### 3.3.4. Case IV: Full model with multiple linear regression fitting

Using the scaling as in Eqs. 2a one can reduce the set of Eqs. 3 into the following second order nonlinear differential equation.

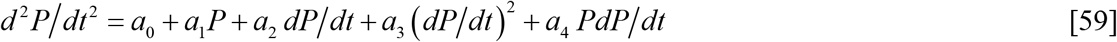

Here *P* is the normalized product concentration and various coefficients multiplying the time derivatives are defined as follows.

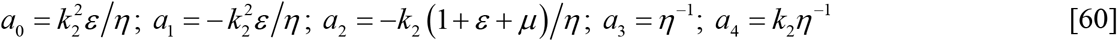

Clearly Eq. 59 can be written as a multiple linear regression model as 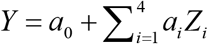 where *Y* and *Z*_1_,…*Z*_4_ are defined as follows.

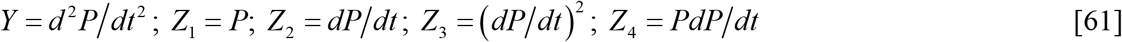

Using the normalized product evolution data (*P, t*) as the input for analysis, one needs to first compute *Y* and *Z*_1_,…*Z*_4_ by numerical differentiation. Then a multiple linear regression fitting procedure needs to carried out to obtain the best fit values (*â_s_*) of various parameters defined in Eqs. 60 along with their standard errors at a given confidence level (*θ* ~ 0.95 in the current scenario) as follows.

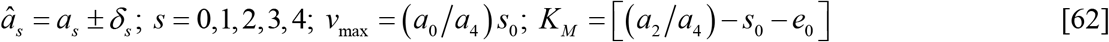

Upon following the error propagation theory one finds the following expressions for the best fit values of steady state MMS enzyme parameters.

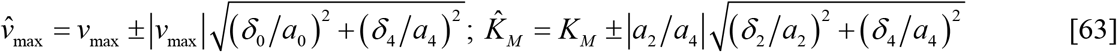

The main advantage of Case IV will be the usage of simple multiple linear least square fitting procedure which yield the best fit parameters in a single step. However this method will not be an efficient one especially when there is a significant amount of noise in the input data. Because fluctuations present in the input product evolution data (*P, t*) will be tremendously amplified upon numerically computing the first and second derivative terms of *P* with respect to time as in Eqs. 61 which may make the best fit parameters meaningless.

#### 3.3.5. Performance of various progress curve models

The relative efficiencies of Cases I, II, III and IV in attaining various MMS parameters from the experimental data are shown in Figs. 10A-F. Detailed analysis clearly suggest that reliable estimates of MMS parameters can be obtained only from the full model fitting as in Case I. Full model fit seems to work very well both in the presence and absence of noise in the input data (Tables 3A and 3B). Integrated form of sQSSA that is given by Case II (S) seems to overestimate both *K_M_* and *v*_max_. On the other hand *φ*-sQSSA as in Case II (P) and multiple linear regression model fit as in Case IV seems to underestimate the steady state MMS parameters in the presence of noise though estimates from multiple linear regression model given in Case IV are exact in the absence of noise in the input data. Nonlinear least square fitting with double exponential function as in Case III seems to work very well in obtaining various timescale components associated with the *φ*-approximation especially 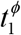, 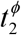 and the timescale separation ratio (Tables 3A and 3B). However the steady state timescale 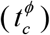 of MMS obtained from double exponential fitting seems to be inaccurate and one always obtains 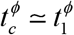 which is evident from the nonlinear least square fitting results. Figs. 4A-D suggest that accurate estimate of 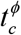 will be possible only when *η* > 1 or *κ* > 10 and the error associated with the φ-approximation of 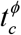 will be maximum when *ε* ~ 1.

**FIGURE 10:**
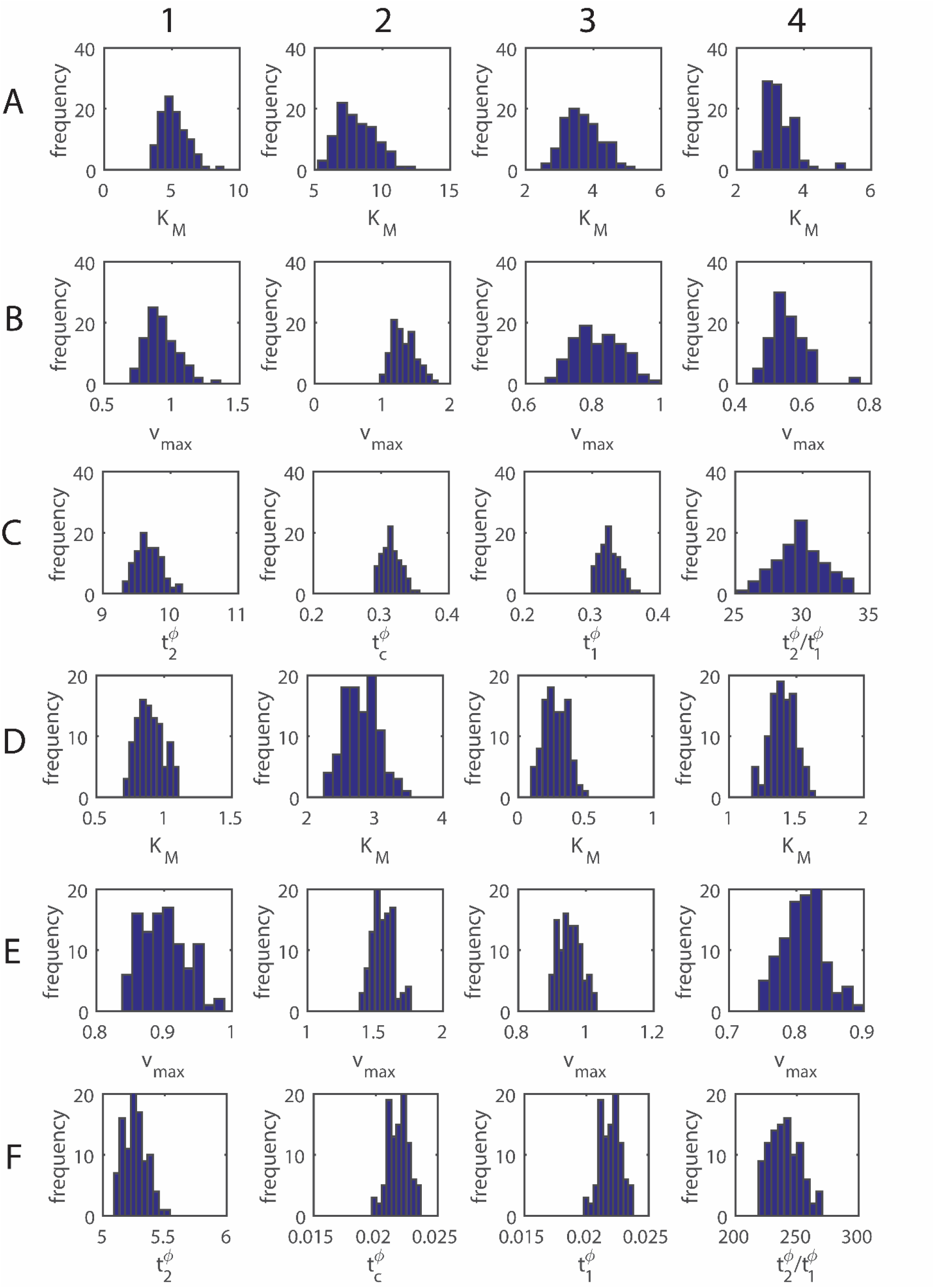
Errors in the progress curve analysis. Sample data were generated using Eqs. 47 with the common settings as *κ* ~ 0.1, *ε* ~ 0.1, *s*_0_ ~ 4.5 mM (so that *e_0_* ~ 0.45 mM) and *γ_X_* = *γ_P_* ~ 10^−3^, *k_2_* ~ 2s^−1^ and *Δt* ~ 10^−2^ s. All the histograms were generated as follows. Totally 100 independent stochastic trajectories were generated by numerically integrating Eqs. 47. Subsequently we used these datasets on (*P, X, S, t*) as input for the nonlinear least square fitting procedures associated with Cases I, II and III using Marquardt-Levenberg algorithm. Histograms were generated using these obtained values of *K_M_, v*_max_ and various timescale components (Case III (X), rows **C** and **F**) at *θ* = 0.95 confidence level from each set of stochastic trajectories. R^2^ in all these fittings were > 0.99. Original parameter values and parameter estimates using various model fitting over noise free datasets i.e. *γ_X_* = *γ_P_* = 0 are given in Table 3A and 3B. **A-C**: *η* ~ 1.0 and one finds the true values of *K_M_* = (*η* + *κ*) *s*_0_ ~ 4.95 mM and *v*_max_ = *k_2εs0_* ~ 0.9 mMs^−1^ as the pre-set values of steady state MMS parameters. **D-F**: *η* ~ 0.1 and one finds the original values of *K_M_* ~ 0.9 mM and *v*_max_ ~ 0.9 mMs^−1^. **Columns** for **A, B, D, E. 1.** Case I, a full model fitting as in Eq. 48. 2. Case II (S), as in Eq. 51 for substrate depletion data. **3**. Case II (P) as in Eq. 53 for product evolution data. **4**. Case IV (P) multiple linear regression fit as in Eq. 61 for product evolution data. Results obtained from full model (Case I) fitting over noisy dataset fairly agree with the original values of MMS parameters. Case II (S) overestimates and Case II (P) underestimates both *K_M_* and *v*_max_. Case IV (P) with multiple linear regression fitting seems to underestimate or overestimate the MMS parameters depending on the value of *φ*. **Columns** for **C** and **F**. double exponential fit with Eq. 57 on the normalized enzyme-substrate complex (*X, t*) dataset. 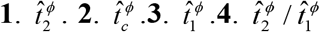, time scale separation ratio.

**Table 3A.** Nonlinear least square fitting results over noise free dataset I.

| Method | $\hat{K}_M$ (mM) | $\hat{v}_{\max}$ (mMs<sup>-1</sup>) | $\hat{t}_1^\phi$ (s) | $\hat{t}_c^\phi$ (s) | $\hat{t}_2^\phi$ (s) |
| --- | --- | --- | --- | --- | --- |
| Case I (P) | $4.95 \pm 2 \times 10^{-5}$ | $0.9 \pm 0.0003$ | - | - | - |
| Case I (X) | $4.95 \pm 3 \times 10^{-8}$ | $0.9 \pm 1 \times 10^{-6}$ | - | - | - |
| Case I (S) | $4.95 \pm 6 \times 10^{-8}$ | $0.9 \pm 4 \times 10^{-7}$ | - | - | - |
| Case II (S) | $7.80 \pm 0.082$ | $1.14 \pm 0.005$ | - | - | - |
| Case II (P) | $3.59 \pm 0.032$ | $0.75 \pm 0.002$ | - | - | - |
| Case III (X) | - | - | $0.32 \pm 0.008$ | $0.33 \pm 0.044$ | $9.62 \pm 0.014$ |
| Case IV (P) | $5.04 \pm 0.0002$ | $0.90 \pm 3 \times 10^{-5}$ | - | - | - |
| Original | 4.95 | 0.9 | 0.232 | 0.91 | 10.76 |

**Note:** settings used to generate datasets are $\eta \sim 1$ , $\varepsilon \sim 0.1$ , $\kappa \sim 0.1$ , $s_0 \sim 4.5$ mM, $e_0 \sim 0.45$ mM, $k_2 \sim 2\text{s}^{-1}$ , $K_M \sim 4.95$ mM, and $v_{\max} \sim 0.9$ mMs<sup>-1</sup>.

**Table 3B.** Nonlinear least square fitting results over noise free dataset II.

| Method | $\hat{K}_M$ (mM) | $\hat{v}_{\max}$ (mMs<sup>-1</sup>) | $\hat{t}_1^\phi$ (s) | $\hat{t}_c^\phi$ (s) | $\hat{t}_2^\phi$ (s) |
| --- | --- | --- | --- | --- | --- |
| Case I (P) | $0.9 \pm 8 \times 10^{-10}$ | $0.9 \pm 1 \times 10^{-8}$ | - | - | - |
| Case I (X) | $0.9 \pm 3 \times 10^{-7}$ | $0.9 \pm 1 \times 10^{-6}$ | - | - | - |
| Case I (S) | $0.9 \pm 6 \times 10^{-9}$ | $0.9 \pm 4 \times 10^{-6}$ | - | - | - |
| Case II (S) | $2.75 \pm 0.031$ | $1.54 \pm 0.009$ | - | - | - |
| Case II (P) | $0.27 \pm 0.032$ | $0.95 \pm 0.0005$ | - | - | - |
| Case III (X) | - | - | $0.0230 \pm 0.009$ | $0.0231 \pm 0.009$ | $5.19 \pm 0.014$ |
| Case IV (P) | $0.95 \pm 1 \times 10^{-12}$ | $0.90 \pm 1 \times 10^{-13}$ | - | - | - |
| Original | 0.9 | 0.9 | 0.038 | 0.195 | 6.45 |

**Note:** settings used to generate datasets are $\eta \sim 0.1$ , $\varepsilon \sim 0.1$ , $\kappa \sim 0.1$ , $s_0 \sim 4.5$ mM, $e_0 \sim 0.45$ mM, $k_2 \sim 2\text{s}^{-1}$ , $K_M \sim 0.9$ mM, and $v_{\max} \sim 0.9$ mMs<sup>-1</sup>.

### 3.4. Dynamical efficiency of single substrate MMS enzymes

The overall dynamical trajectories of single substrate MM enzymes over VPS phase space start at (*V, P, S*) = (0, 0, 1) at *τ* = 0 and end at (*V, P, S*) = (0, 1, 0) at *τ* → ∞ with the condition that all the originating trajectories should lie on the plane defined by *V* + *P* + *S* = 1. The length of the curve (we denote it as *L_A_*) which connects these start and end points of MMS dynamics over VPS space can be expressed as follows.

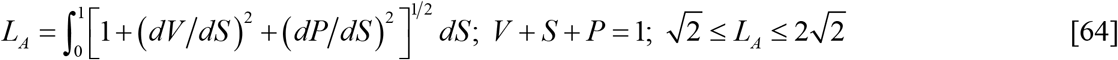

In this equation [*dV*/*dS*] and [*dP*/*dS*] are defined as in Eqs. 20 and 30 respectively. One needs to first integrate these nonlinear ODEs corresponding to *V* and *P* and express them in terms of *S*. Subsequently upon substituting *V* and *P* as functions of *S*, one can numerically evaluate the integral in Eqs. 64 with respect to *S*. Clearly the shortest path between the initial and final points of MMS in VPS space will be via the line defined by *P* + *S* = 1 of the PS plane where *V* = 0 and one finds that 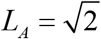 (Figs. 11A1-4). This is the line defined by the intersection of the planes defined by *V* = 0 and *V* + *P* + *S* = 1. The longest path length of MMS dynamics occurs when *ε* → ∞ where it will be partially via the line *V* + *S* = 1 of VS space [i.e. from (*V, P, S*) = (0, 0, 1) to (*V, P, S*) = (1, 0, 0)] and then via the line *V* + *P* = 1 of VP space [i.e. from (*V, P, S*) = (1, 0, 0) to (*V, P, S*) = (0, 1, 0)]. In the trajectory with the longest path length, the steady state will be defined by (*V_c_, P_c_, S_c_*) = (1, 0, 0) which occurs at *τ* = *τ_c_*. Upon substituting the expressions for [*dV*/*dS*] and [*dP*/*dS*] in Eqs. 64 one obtains the following formula for the reaction path length of MMS enzymes for finite values of *ε*.

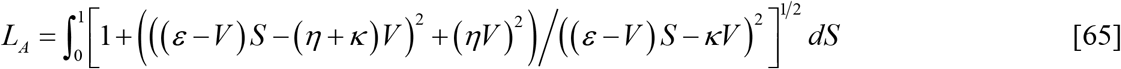

**FIGURE 11:**
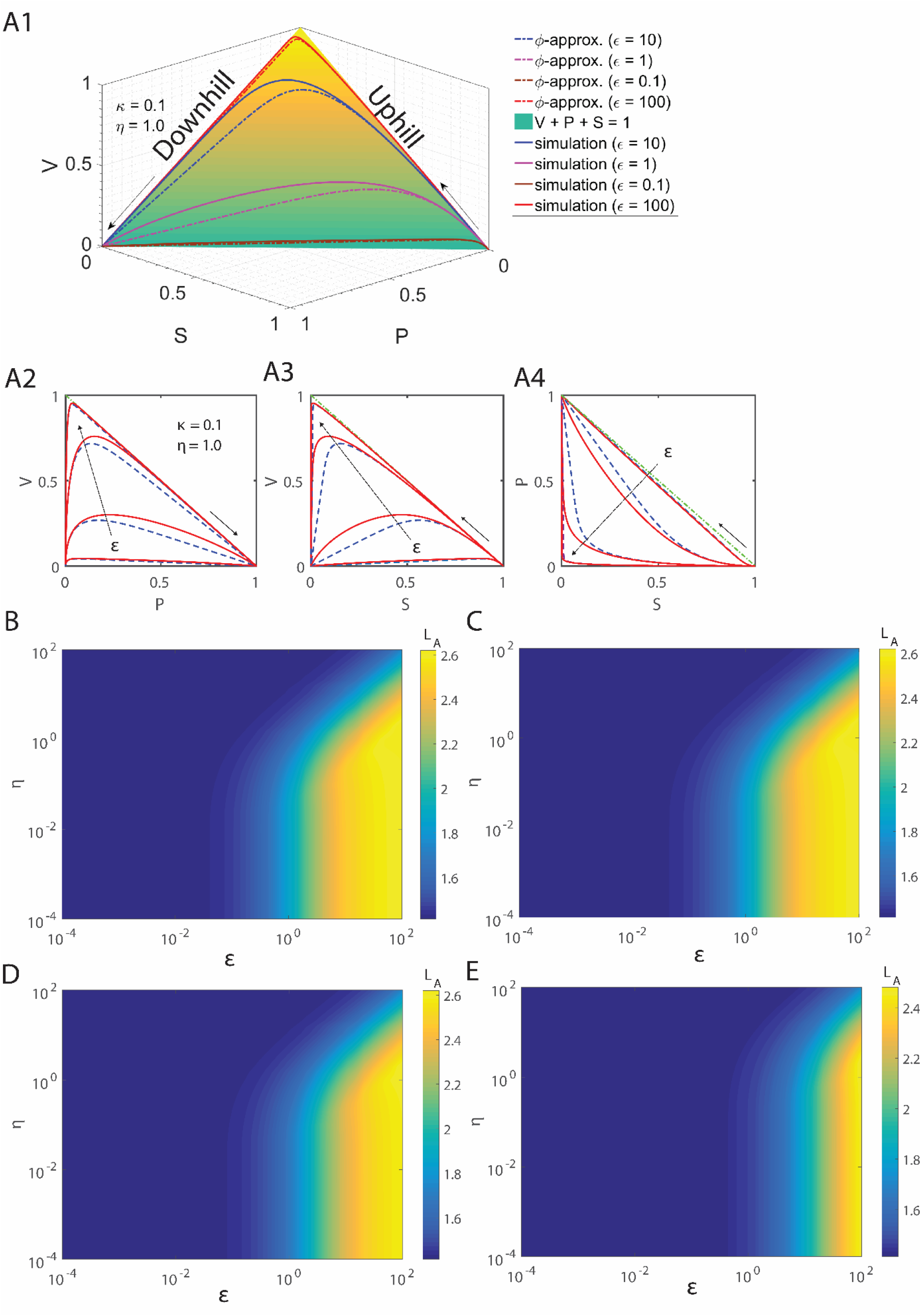
**A1**. Arc or path lengths (*L_A_*) associated with the trajectories of MMS in VPS space. Here the trajectory starts from (*V, P, S*) = (0, 0, 1) at *τ* = 0 and ends at (*V, P, S*) = (0, 1, 0) at *τ* → ∞ with the condition that all the emanating trajectories should lie on the plane defined by *V* + *P* + *S* = 1. Clearly the shortest path between these two points will be via the line defined by *P* + *S* = 1 of the PS plane where *V* = 0 and obviously the shortest path length is 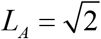. Since *V* = 0 throughout the trajectory here, this shortest route will take infinite amount of time. The longest route will be partially via *V* + *S* = 1 of VS space (uphill) and then via *V* + *P* = 1 of VP space (downhill). The total length of this longest route will be 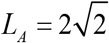. The total path lengths of all those trajectories will lie inside 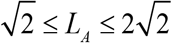. By changing the values of *ε* one can choose the desired reaction path where the path length is positively correlated with *ε*. **A2**. Path lengths 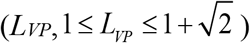 associated with the trajectories of MMS in VP space. **A3**. Path lengths 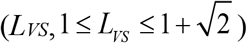 associated with the trajectories of MMS in VS space. **A4**. Path lengths 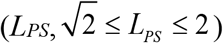 associated with the trajectories of MMS in PS space. In **A2-A4** the direction of dotted arrows indicates iteration over *ε* in (0.1, 1, 10, and 100). Here solid red lines are the simulated trajectories and dotted blue lines are the φ-approximations. Green solid lines are the equations V + P = 1 (**A2**), V + S = 1 (**A3**) and P + S = 1 (**A4**). **B-E**. Variation of the total path lengths of MMS trajectories with respect to changes in the parameters (*ε, η, κ*). **B**. *κ* ~ 10^−2^. **C**. *κ* ~ 10^−1^. **D**. *κ* ~ 1. **E**. *κ* ~ 10.

This integral is not expressible in terms of elementary functions. Further Eq. 65 suggests that *L_A_* is strongly dependent on the parameters (*ε, η, κ*) and one can directly obtain the following limiting conditions (Figs. 11B-E).

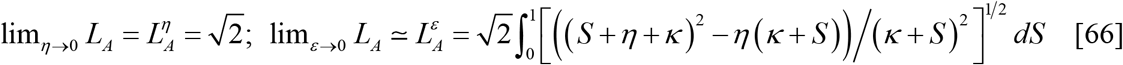

Here one also should note that 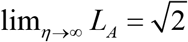 and the maximum of *L_A_* occurs at *η* = 1 (Figs.11B-E). This is a reasonable observation since the maximum possible value of *V* occurs at this point (*ε* and *κ* are fixed) which is positively correlated with *L_A_*. The reaction path lengths associated with the MMS enzymes in various reduced dimensional spaces can be expressed as follows (Figs. 11A2-4).

*Reaction path length in VS space:*

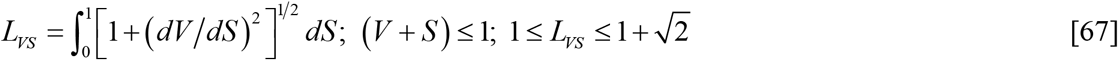

Upon substituting the definition of [*dV*/*dS*] from Eq. 20 in to this equation one finds the following limiting conditions (Fig. 11A2).

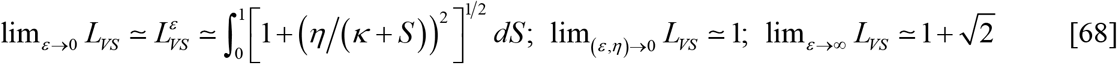

*Reaction path length in VP space:* (Fig. 11A3)

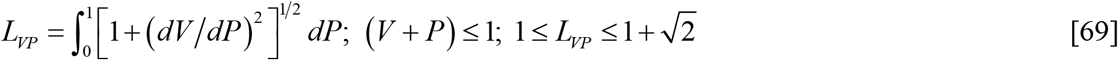

*Reaction path length in PS space:* (Fig. 11A4)

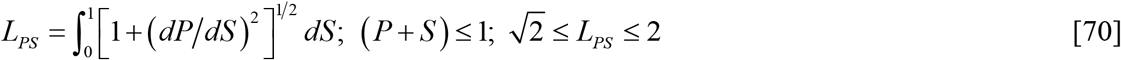

The following limiting conditions exists for the reaction path lengths associated with the dynamics of MMS enzymes in the VP and PS spaces (Figs. 11A2-4).

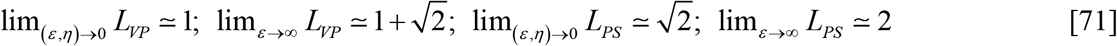

As we have shown in the theory section 2.2, the **catalytic efficiency** of a MMS enzyme will be inversely proportional to the overall average reaction time *τ_T_*. Since exact expression for *τ_T_* is unknown, we use its corresponding *φ*-approximation 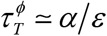 here. Figs. 12A-B show how this total reaction time vary with respect to the control parameters (*ε, η, κ*). In the limit as *ε* → ∞ one finds that 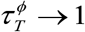 i.e. MMS will behave as a first order decay with an overall rate constant equals to *k_2_*. The overall **dynamical efficiency** of a single substrate MMS enzyme can be measured by the ratio between the total path length *L_A_* in the VPS phase space and the average reaction time required by the MMS enzyme system to travel from (*V, P, S*) = (0, 0, 1) to (*V, P, S*) = (0, 1, 0) in the VPS phase space as follows (Figs. 12E-H).

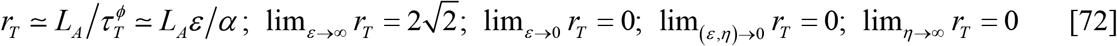

**FIGURE 12:**
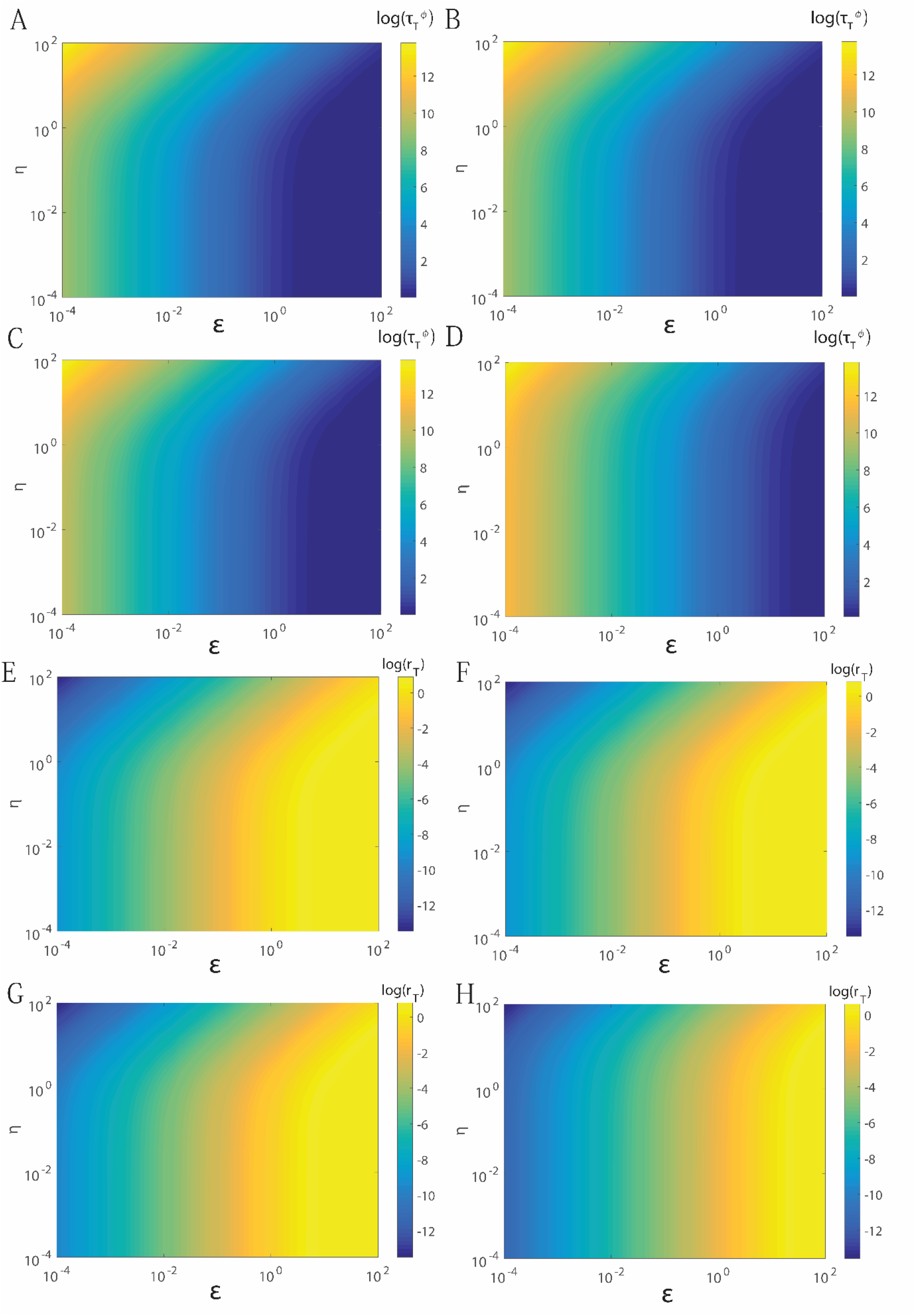
**A-D**. Average reaction time (*τ_T_*) that is required by the MMS enzyme to convert all the initial substrate molecules into product as predicted by *φ*-approximation 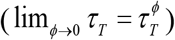. From Eqs. 8 one obtains 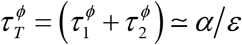. In the limit as *ε* → ∞ one finds that 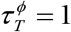 i.e. MMS will behave as a first order decay with an overall rate constant equal to *k_2_*. Variation of 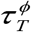 with respect to changes in the parameters (*ε, η, κ*). **A**. *κ* ~ 10^−2^. **B**. *κ* ~ 10^−1^. **C**. *κ* ~ 1. **D**. *κ* ~ 10. **E-H**. Variation of the ratio 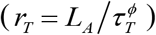 between the reaction path lengths (*L_A_*) associated with the trajectories of MMS in VPS space and the average reaction time (*τ_T_*) that is required by the MMS enzymes to convert the entire initial substrate into product as predicted by the *φ*-approximation 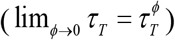. From Eqs. 8 one obtains 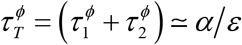. In the limit as *ε* → ∞ one finds that 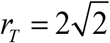 i.e. MMS will behave as a typical first order decay process with an overall rate constant equal to *k_2_*. Variation of *r_T_* with respect to changes in the parameters (*ε, η, κ*). **E**. *κ* ~ 10^−2^. **F**. *κ* ~ 10^−1^. **G**. *κ* ~ 1. **H**. *κ* ~ 10.

Here the dynamical efficiency can be used to measure the fastness of an enzyme in clearing the substrate from the reaction medium. Figs. 12E-H show how the dynamical efficiency *r_T_* of single substrate MMS enzymes varies with respect to the parameters (*ε, η, κ*).

## 4. Conclusions

Analytical solution to the rate equations associated with the single substrate MMS enzymes is not known. Several groups across various fields tried to provide approximate formulas which can be used along with other experimental techniques to obtain various kinetic and steady state parameters of a given MMS enzyme. The quasi steady state approximation (QSSA) with stationary reactant assumption is the widely adopted one which is applicable mainly to the post-steady state dynamics of MMS enzymes. Depending on the relative initial concentrations of enzyme and substrate one can choose among tQSSA, sQSSA and rQSSA (**t**otal, **s**tandard and **r**everse). Here sQSSA will be applicable when the initial substrate concentration is much higher than the concentration of enzyme. When the initial enzyme concentration is much higher than substrate then rQSSA will be applicable. It seems that tQSSA is applicable under both the conditions to certain extent. In this article we have reformulated the rate theory associated with the dynamics of single substrate MMS enzymes.

Here we have introduced yet another scaling scheme for the rate equations associated with the MMS dynamics. We identified the critical parameters which can completely characterize the entire dynamics of single substrate MMS enzymes and identified the validity range of various QSSA methods in line with these parameters. We reformulated the rate equations of MMS over velocity-substrate, velocity-product, substrate-product and velocity-substrate-product spaces and obtained various approximations for both pre- and post-steady state dynamics of MMS. Using this framework under certain limiting conditions we could successfully compute the steady state, pre- and post-steady state timescales associated with the dynamics of MMS enzymes. We have also computed the approximate values of steady state velocity, substrate and product. We further redefined the catalytic efficiency and defined the dynamical efficiency of MMS enzymes as the ratio between the reaction path length of MMS enzymes in the velocity-substrate-product space and the average total reaction time. Here dynamical efficiency completely characterizes the phase-space dynamics of MMS enzymes and it would tell us how fast the MM enzyme can effectively clear a harmful substrate from the environment. We subsequently performed a detailed error level analysis over various approximations along with the already existing QSSA and progress curve models. Finally we have discussed the positive and negative points corresponding to various steady state and progress curve models.

